# Environmental heterogeneity drives tsetse fly population dynamics and control

**DOI:** 10.1101/493650

**Authors:** Hélène Cecilia, Sandie Arnoux, Sébastien Picault, Ahmadou Dicko, Momar Talla Seck, Baba Sall, Mireille Bassène, Marc Vreysen, Soumaïla Pagabeleguem, Augustin Bancé, Jérémy Bouyer, Pauline Ezanno

## Abstract

A spatially and temporally heterogeneous environment may lead to unexpected population dynamics. Knowledge still is needed on which of the local environment properties favour population maintenance at larger scale. For pathogen vectors, such as tsetse flies transmitting human and animal African trypanosomosis, such a knowledge is crucial to design relevant management strategies. We developed an original mechanistic spatio-temporal model of tsetse fly population dynamics, accounting for combined effects of spatial complexity, density-dependence, and temperature on the age-structured population, and parametrized with field and laboratory data. We confirmed the strong impact of temperature and adult mortality on tsetse populations. We showed that the coldest cells with the smallest variations in temperature acted as refuges when adult mortality was homogeneously increased, control being less effective in such refuges. In contrast, optimizing control by targeting the cells contributing the most to population management resulted in a decline in population size with a similar efficacy, but resulted in more dispersed individuals, control efficacy being no longer related to temperature. Population resurgence after control was slow, but could be very high locally in refuges. Situations were highly contrasted after a heterogeneous control, refuges being located at the interface between controlled and uncontrolled zones. Our results highlighted the importance of baseline data collection to characterize the targeted ecosystem before any control measure is implemented.

## Introduction

Environmental spatial heterogeneity is an important driver of insect population dynamics (Tilman & Kareiva 1997; Vinatier et al. 2011), inducing insect movements from source to sink patches and possibly enhancing population persistence in unsuitable patches (Holt 1985; Pulliam 1988). In addition, environmental suitability varies over time both at local scale, due to microclimate variations as related to vegetation growth (Keppel et al. 2017), and at a larger scale, due to seasonal occurrence of unfavourable periods. Confounding the role of spatial and temporal environmental heterogeneity can potentially result in erroneous predictions of ecological processes (Clark 2005). However, relating such a complex time- and space-varying habitat with population dynamics still is a challenge in ecology (Sutherland et al. 2013; Crone 2016; Griffith et al. 2016). Therefore, examples of the complex interplay between spatio-temporal environmental variability and population dynamics can illustrate theoretical concepts and assess which patch properties (co)contribute to define sources and sinks in heterogeneous environments.

This is particularly true when it comes to managing vector-borne diseases whose transmission may be affected by landscape configuration as interactions between hosts and vectors largely depend on their and temporal variations in environmental suitability could induce unexpected changes in the dynamics of the vector population. Despite this, insect pest management strategies are often designed and implemented without considering local environmental specificities, potentially reducing the chances of success.

Tsetse flies (*Glossina* spp.) are vectors of African trypanosomes, widely recognized as a major pathological constraint for productive livestock and for sustainable agricultural development in sub-Saharan Africa (Alsan 2015). *Trypanosoma* spp. parasites cause both Human African Trypanosomosis (HAT, sleeping sickness) in humans and African Animal Trypanosomosis (AAT, nagana) in livestock. Tsetse flies are widely distributed in Africa and occur in 38 countries infesting around 10 million km^2^ (Vreysen et al. 2013). Over 60 million people are continuously exposed to the risk of becoming infected with HAT, a neurological, potentially lethal disease, mainly in remote rural areas where access to health services is very limited. In addition, farmers in tsetse-infested areas suffer up to 20-40% losses in livestock productivity which amounts to an estimated annual loss of $4,500 million (Budd 1999). Although there are 31 species and subspecies of tsetse flies described, only a third is of economic (agricultural and veterinary) and human health importance (Solano et al. 2010a). Efforts to manage the vector and the disease in Africa have been on-going for decades but have largely failed to create sustainable tsetse-free areas, and it is estimated that the tsetse distribution has only been reduced with less than 2% (Allsopp 2001; Bouyer et al. 2013a). Although the ecology and biology of tsetse flies are rather complex, their very low rate of reproduction (one offspring every 10 days) make them an ideal target for eradication strategies, but this would require a better understanding of their spatio-temporal dynamics (Peck & Bouyer 2012).

Mathematical models have proved to be relevant tools in insect ecology, to better understand the dynamics of insect populations (Hasting 2012) and to predict these dynamics under changing conditions (Evans et al. 2012). Process-based models incorporate at minimal costs sparse and heterogeneous knowledge from various areas, species, and fields of expertise. Simulations are complementary to field observations and experiments (Restif et al. 2012), enabling the fast acquisition of quantitative predictions which can in turn emphasize the need for further biological investigations. Moreover, the range of behaviours of complex systems can be scanned using mechanistic models, and scenarios can be tested (Cailly et al. 2012). Provided hypotheses and limits are clearly stated (Getz et al. 2018), models can guide decision-making (Sutherland & Freckleton 2012).

With respect to tsetse biology and population dynamics, entomologists have developed a number of models (Rogers 1988, 1990; and more recently: Vale & Torr 2005; Lin et al. 2015), and encouraged their use when making management decisions (Hargrove 2003; Childs 2011; Meyer et al. 2018). However, most models have failed to predict the persistence of target populations leading to inaccurate guidelines for control programs (Peck & Bouyer 2012; Bouyer et al. 2013b). In addition, most of these programs were not implemented following area-wide principles (Klassen 2005; Hendrichs et al. 2007) and their failure could be due to population resurgence in non-eradicated patches or re-invasion of the targeted zone by neighbouring populations (Meyer et al. 2016; Lord et al. 2017). With respect to tsetse flies, it is still unclear what defines relevant patch properties and how they define sources and sinks in a hostile environment created by eradication efforts. Spatial complexity of the environment has been shown to considerably influence model predictions (Peck 2012; Barclay & Vreysen 2013; Lord et al. 2017), and population dynamics will be different amongst local patches of variable suitability, possibly affecting population dynamics at the larger metapopulation scale.

Our objective was to assess the effect of spatial and temporal heterogeneity of the environment on the dynamics of tsetse fly populations at the metapopulation scale, as well as the effects of spatially targeted treatments on adult fly mortality and hence, on fly population densities. We have developed an original mechanistic spatio-temporal model of tsetse fly population dynamics that incorporates environmental heterogeneity through a data-driven approach. The model was applied to *Glossina palpalis gambiensis* population of the Niayes (Senegal), a region with an ongoing eradication project (Dicko et al. 2014; Vreyssen et al. in press). In this area, less than 4% of the habitat was considered favourable for *G. p. gambiensis* (Bouyer et al. 2010), and the tsetse populations were highly structured across the metapopulation (Solano et al. 2010b). This knowledge was incorporated in the model, accounting for combined effects of spatial complexity, density-dependence, and temperature on the age-structured population.

## Material and methods

### Key knowledge on tsetse biology

Meteorological variables influence the abundance and spatio-temporal distribution of arthropod disease vectors (Hay et al. 1996). Effect magnitude depends on species (Rogers & Randolph 1991; Rogers et al. 1996; Hargrove 2001), but for tsetse flies, average temperature is the most influential meteorological variable on life cycle (Hargrove 2004). However, its influence compared to, or combined with, demographic processes is poorly understood.

Tsetse flies reproduce by adenotrophic viviparity, i.e. the egg hatches in the female’s uterus and the larva is nourished by the milk glands until larviposition (Fig. 1A). Between a temperature range of 20-30°C, decreasing temperatures will increase the period between larvipositions (Harley 1968). Similarly, colder temperatures will prolong the pupal period (Glasgow 1963; Phelps & Burrows 1969a,b). The newly emerged fly (called thereafter nulliparous female up to her first larviposition) takes its first blood meal to fully develop its flight muscles and reproduce. Depending on species and temperature, maturation of the first oocyte in the female fly takes about 18 days, making the period between emergence and first larviposition longer than between subsequent larviposition events (10 days, Hargrove 2004).

**Figure 1.**
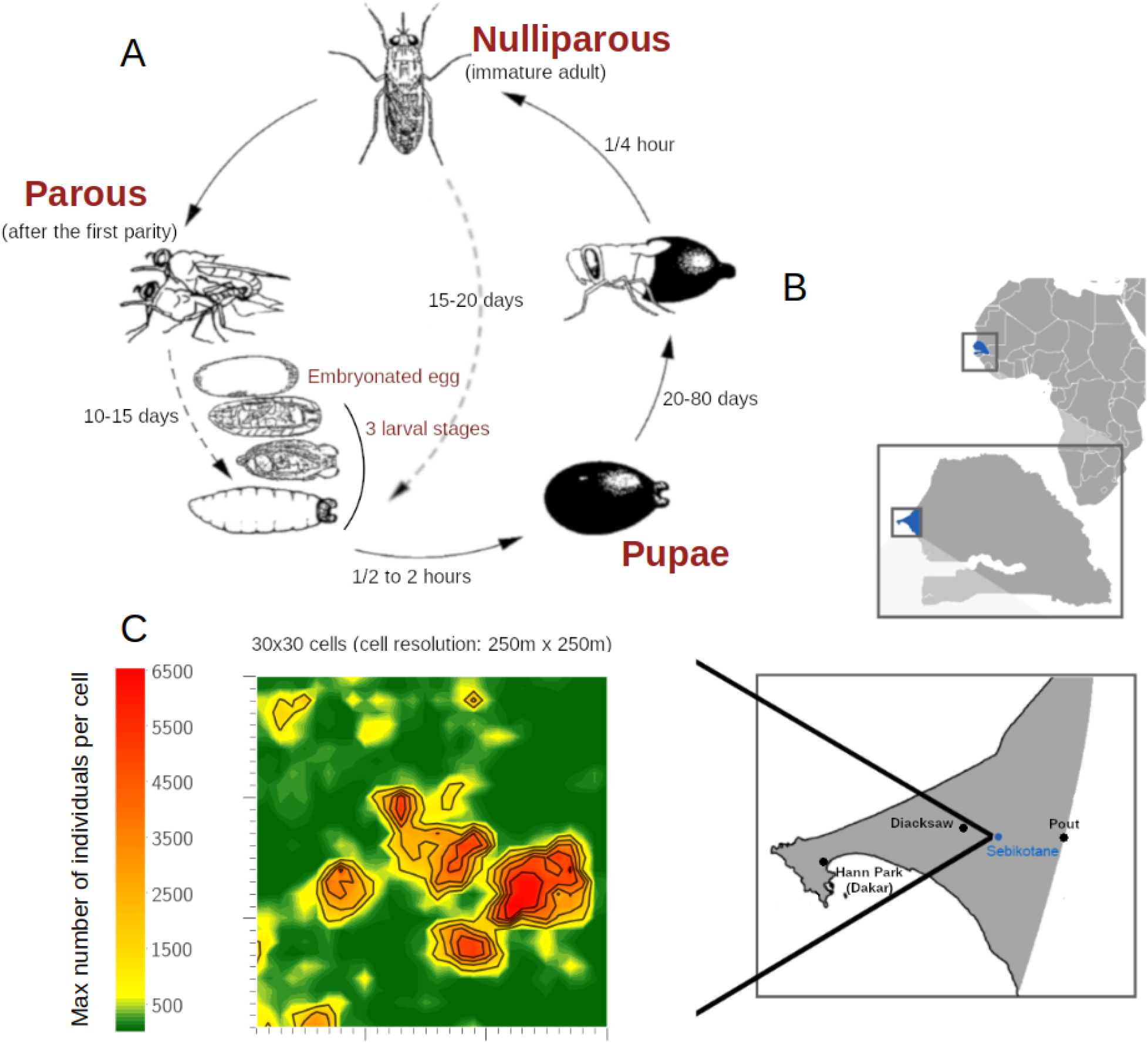
Local and general tsetse fly population dynamics applied to the Niayes in Senegal. (A) life cycle of tsetse flies as it occurs within each cell (drawn by D. Cuisance). (B) Map of Senegal with locations where field data were collected. (C) Simulated area, highlighting the spatial heterogeneity resulting in different local carrying capacities *k*_*c*_.

Extreme cold (below 20°c) or warm (upon 30°c) temperatures increase fly mortality (Hargrove 2001). Mortality, related to predation and feeding success, is density-dependent (Rogers & Randolph 1984) and age-dependent (Hargrove 1990), with remarkably high losses in nulliparous flies partly due to starvation risk (Phelps & Clarke 1974; Hargrove 2004). Learning capability of older flies makes them return on their first host, which increases their hunting efficiency with age (Bouyer et al. 2007).

Tsetse flies are classified into three groups based on their behaviour, habitat preference and distributions, i.e. forest (subgenus *Fusca*), savannah (subgenus *Morsitans*), and riverine flies (subgenus *Palpalis*). Most of previous models concerned the savannah species *Glossina pallidipes* and *G. morsitans*. We focused on the riverine species *G. p. gambiensis* which thrives in forest galleries and riparian thickets (Bouyer et al. 2005). The habitat of this species stretches along rivers and therefore, its dispersal is mostly in one dimension. In some areas, like the Niayes of Senegal (Fig. 1B), climate changes have resulted in rivers and associated vegetation disappearing, and *G. p. gambiensis* consequently adapted to patchy vegetation mainly associated with human irrigation activities (Bouyer et al. 2010) and disperse in two dimensions like tsetse flies of the *fusca* and *morsitans* groups. Furthermore, isolated populations in fragmented habitats are ideal targets for eradication strategies within area-wide integrated pest management approaches (Hendrichs et al. 2007; Bouyer et al. 2015). Hence, our case study is of broad relevance for better understanding and predicting tsetse fly spatio-temporal population dynamics in rapidly changing ecosystems that are gradually becoming the norm (Guerrini et al. 2008).

### Data on tsetse biology

The effect of temperature on mortality and fecundity of *G. p. gambiensis* was assessed under experimental conditions (Pagabeleguem et al. 2016). We used data of the first larviposition period (time between emergence and first larviposition day) and of subsequent inter-larval periods (time between reproductive cycles). The colony was maintained at 24°C and only temperatures above 24°C were assessed in terms of the maximum critical temperature for the flies. Therefore, most data used to estimate female mortality were obtained at 24°C and none at lower temperatures. In addition, the effect of temperature on the length of the pupal period was measured under experimental conditions at the Centre International de Recherche-Développement sur l’Elevage en zones Subhumides (CIRDES) in Bobo-Dioulasso, Burkina Faso, in 2009. One-hundred-and-twenty-day-old pupae were held in climate controlled rooms until emergence. The experiment was replicated three times for each temperature tested (Table S1).

Dispersal of *G. p. gambiensis* was assessed from release-recapture data of marked sterile males from October 2010 to December 2012 (Pagabeleguem 2012). Mass-reared male flies from the CIRDES colony were shipped as irradiated and chilled pupae to Senegal (Pagabeleguem et al. 2015) and after emergence, released twice a month and monitored in four areas: Parc de Hann in Dakar, Diacksaw Peul, Pout, and Kayar (Fig. 1B). Two release points were selected per location (in suitable vs. unsuitable habitats) and released flies were trapped using Vavoua traps (Laveissière & Grébaut 1990) that were deployed at intervals of 100-300 m up to 2 km from the release points. Traps were deployed before 9 am and collected after 4 pm 3 days later. The monitoring of a release stopped when less than 2 marked males were recaptured.

In another study, natural abortion rates were monitored in the Parc de Hann, Diacksaw Peul, Sebikotane, and Pout (Fig. 1B). In each site, 10 traps were deployed for three days every month from March 2008 to February 2009, and then every three months until September 2010 (Hann, Diacksaw) or December 2011 (Pout, Sebikotane). Flies were collected at least once a day and female flies were dissected to assess their ovarian age. This female dataset was used to calculate the population age structure, to be compared to simulation results for partial validation of the model.

### Environmental data

The spatio-temporal heterogeneity of the environment was realistically represented using an original data-driven approach. The environmental carrying capacity and the local daily temperatures were incorporated in the model.

The carrying capacity was defined as the maximum sustainable number of individuals for a given area and was estimated as (Eq. 1, Fig. 1C):

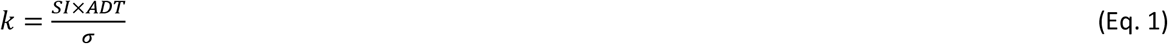

where *SI* is the suitability index as estimated with a species distribution model (Dicko et al. 2014) that was based on maximum entropy (Maxent) (Supporting Information 2.1), *σ* is the trap efficiency, i.e. the probability that a trap catches a fly within 1 km^2^ within a day (Barclay and Hargrove 2005), and *ADT* is the apparent density of the fly population (no. flies per trap per day, Dicko et al. 2015). All available trap catch data collected in the Niayes before the start of the eradication campaign (2007-2010) were used to estimate local carrying capacities (Supporting Information 2.1).

Air temperatures measured in weather stations are not those experienced by flies in resting places. Tsetse flies prefer micro-environments that are normally 2-6°C cooler than the ambient temperature (Hargrove & Coates 1990). In addition, temperature generally increases from the centre of a gallery forest towards its edges (Bouyer 2006). Therefore, the micro-climate was modelled by approximating local temperatures truly perceived by the tsetse flies. High resolution macro-climate data were freely available for 2011 in the studied area and were corrected using temperature data recorded in selected suitable habitat patches (Supporting Information 2.2). Approximated temperatures were used as model inputs in a zone known as suitable for tsetse to check if the simulated population persisted as expected.

### A mechanistic spatio-temporal model of tsetse fly population dynamics

A mechanistic and deterministic compartmental model was developed to predict the spatio-temporal dynamics of *G. p. gambiensis* population accounting for environmental heterogeneity and density-dependence. Individuals were categorized into pupae (P), without differentiating males and females, nulliparous females (N), and parous females with four ovarian age categories (F_1_, F_2_, F_3_, F_4+_, Fig. S1, Hargrove & Ackley 2015). Adult males (M) were not considered limiting for breeding. They could mate from the age of day 6 post-emergence, regardless of temperature, after which they were only subject to mortality (Solano et al. 2010a), and they played a role in density-dependent processes. The environment was modelled using a grid (cell resolution: 250m × 250m; study area: 30 × 30 cells; Fig. 1C). The model was implemented in Python as a discrete-time model with a one-day time step (Supporting Information 7). Parameter values are provided in Table S2.

#### Within-cell dynamics

The population size of life stage *S* at time *t* in cell *c* decreased with mortality, following a negative exponential model of instantaneous rate *μ*_*S,t,c*_ (Eq. 2, Table S2). Considering the lack of data on pupa mortality, we used a constant rate (Eq. 3, Table S2, Childs 2011). For adults, the log of mortality rates increased linearly with temperature (θ_*t,c*_ at time *t* in cell *c*) above 24°C (Hargrove 2004). Below this threshold, and for the range of temperatures observed in the field, the literature and the lack of data suggested a constant mortality rate (Eq. 4, Table S2). Age-dependence was featured by setting nulliparous mortality to twice that of parous females (Alderton et al. 2016, Eq. 5). Density-dependence occurred when the adult population exceeded the cell carrying capacity (Eq. 6–7, Table S2, Hargrove 2004).

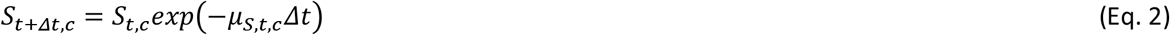

with stage *S* ∈ {*P*, *N*, *F*_*x*_, *M*} and ovarian age *x* ∈ {1, 2, 3, 4+} (note that *μ*_*F,t,c*_ applied irrespective of ovarian age), *Δt* = 1, and:

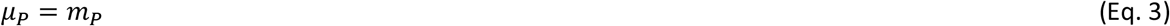

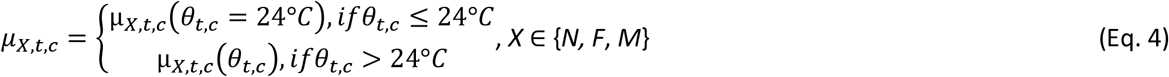

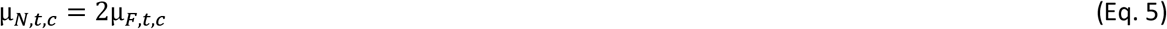

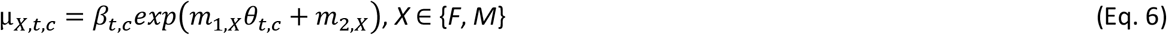

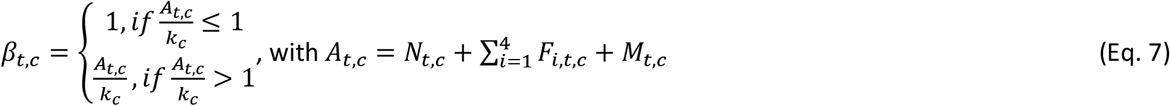

In addition, individuals evolved within and between stages as a function of temperature. Pupa development function *δ*_*P,t,c*_ was fitted to the data. For nulliparous and parous females, consistency of experimental data on the target species was checked against published equations (Hargrove 2004, Eq. 8, Table S2, Fig. S4):

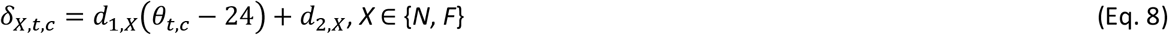

Each stage was discretized into *n*_*S*_ states, *n*_*S*_ being the longest duration in stage *S* obtained with its development rate *δ*_*S*_ calculated at the minimum temperature of the year *min*(*θ*_*t,c*_) (Fig. S1). For higher temperatures, individuals made a leap forward in the development vector, the interval being determined by the integer part *l* of *Δ* (Eq. 9, Fig. S1).

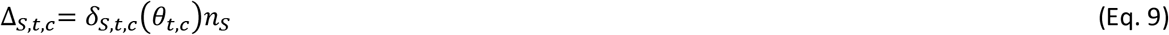

To avoid discretization artefacts, individuals were proportionally divided into two successive states according to the decimal part *q* of Δ (Fig. S1). Individuals who reached state *n*_*S*_ (i.e. stage *S* is completed) evolved to the next stage. A pupa was produced at the end of both nulliparous and parous female stages. After the fourth ovarian age, parous females looped back to the start of F_4+_ (i.e. stage F_4+_ represented females who have produced at least 4 pupae).

#### Between-cell dynamics

An original dispersal pattern was designed favouring suitable over hostile habitats to be conform with species behaviour. The proportion *p*_*t,c*_ of flies leaving cell *c* at time *t* was controlled by a sigmoidal density-dependent dispersal rate (Lloyd-Smith, 2010), adapted for individuals competing to access resources (Rogers & Randolph 1984) (Eq. 10, Table S2):

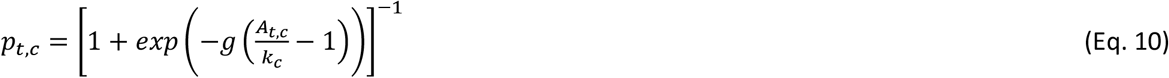

where *k*_*c*_ denotes the carrying capacity in cell *c*, *A*_*t,c*_ denotes the number of adults in cell *c* at time *t*, and *g* denotes a shape parameter set to 10 meaning that 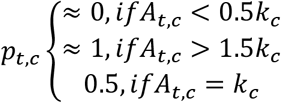 (Fig. S2).

The spatial distribution of dispersing flies from cell *c* to neighbouring cells *Prob*_*c*→*i*∈{*v*}_ was determined by the relative attractiveness of neighbouring cells *a*_*t,i*∈{*v*}_ (Eq. 11–12). This attractiveness was designed to favour the emptiest cells (*A*_*t,i*_ ≪ *k*_*i*_) and cells of greatest *k*_*i*_ in case of equal *A*_*t,i*_/*k*_*i*_. An extended Moore neighbourhood of range *r* was used: flies dispersed from a cell to its (2*r* + 1)^2^ neighbours (*v*), which included the cell itself and diagonals. Parameter *r* is the maximum distance reached daily, in number of cells, rather than the effective distance covered per fly per day, as the trajectory is not linear. It was calibrated using data by taking into account the average 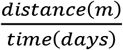 between release and capture of marked flies (Fig. S3).

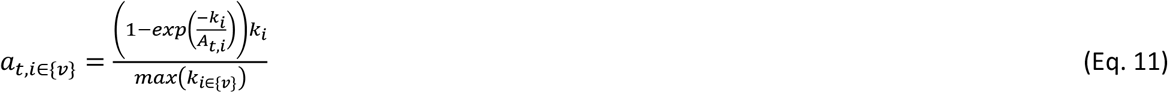

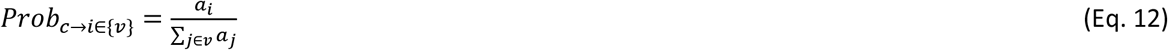

### Model setting and sensitivity analysis

A 3-year burn-in period was simulated starting with N_0,c_=M_0,c_=0.5*k*_*c*_ (A_0,c_=*k*_*c*_), using reference parameter values (Table S2). This provided the initial conditions for the no-control scenario and for the model sensitivity analysis, where population dynamics was simulated over three more years. Carrying capacities were spatially heterogeneous (Fig. 1C) but assumed constant over time. Daily perceived temperatures were estimated per cell for one year and these were repeated during the following years.

The individual and joint effects of input variations on aggregated output variance (Table S3) were evaluated through a variance-based global sensitivity analysis using the Fourier Amplitude Sensitivity Testing (FAST) method (Saltelli et al. 2008). Population size and age structure were outputs of interest. As traps do not catch nulliparous and females of ovarian age “4 and more” as efficiently as females of intermediate ovarian ages (Sanders 1962), predicted age structure was compared with field data for females of ovarian age 1, 2, and 3: 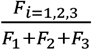. Mortality and development functions of each life stage were varied using multiplying factors (i.e. function formulas were kept). The reference values of multiplying factors were all equal to one. A common factor was applied to all adult mortalities (N, M, F_1:4+_) to maintain a similar order of values. A multiplying factor was also applied to carrying capacities to regulate the magnitude of density-dependence. As the dispersal rate should remain in the range [0-1], the shape parameter *g* was varied (Fig. S2). Parameters and multiplying factors were varied by ± 5% of their reference value. The same range, when applied to temperature, changed the annual mean by more than 2°C, which was far greater than what was observed. Therefore, a variation of ± 0.3°C was used, corresponding to the average deviation from the daily mean in the area (Fig. S6). First order and interaction sensitivity indices were calculated per parameter (Saltelli et al. 2008).

### Evaluation of control strategies

A spatially targeted control strategy was mimicked by increasing adult mortality during one year, starting from the same initial conditions as in the no-control scenario. For successively reduced proportions of controlled cells (starting with 100% of the cells being controlled), we assessed the minimal mortality increase needed to decrease the female population size down to 2 or 5% of its initial size over the whole grid after one year. We stopped reducing the proportion of controlled cells once it became impossible to achieve the targeted population reduction whatever the mortality rate. We defined a score to optimize the selection of controlled cells. The best location of controlled cells was defined by assessing the contribution of each cell *j* to the total female population over the grid (*n* cells) at the end of the control (*t* = 1 year) if cell *j* was not controlled (Eq. 13):

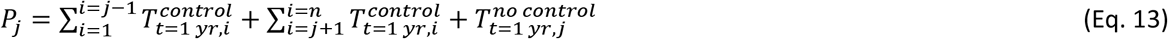

where the total number of females in cell *c* was: *T*_*t*=1 *yr,c*_ = *N*_*t*=1 *yr,c*_ + *F*_1:4+,*t*=1 *yr,c*_.

Cells with the highest *P*_*j*_ were given priority control. As a result, optimized strategies were defined by the minimal proportion of cells to be controlled, their optimal location, and the control effort required.

The control efficacy was assessed with respect to the female population size at both grid and cell scales. We computed for each cell *c* after one year of control: (1) the proportion of females in the area which were located in that cell, 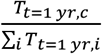, which indicated cells with the highest proportion of the female population; (2) the abundance of females in cell *c* in the control vs. the no-control scenarios after one year 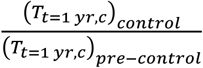, which quantified the local impact of increased mortality.

Then, population resurgence was simulated for one more year after the end of the control period, taking into consideration the reference value of female mortality. To identify the cells that contributed most to the recovery of the population, the local growth rate was calculated per cell: 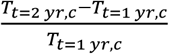, *T*_*t*=2 *yr,c*_ being the female abundance in cell *c* one year after the end of the control period (t = 2 years).

We analysed the relationships between the local environmental variables (carrying capacity, mean temperature, and temperature variance in each cell) and these three cell indicators, reflecting different properties of the population spatial structure.

## Results

### New insights from biological data

New equations were calibrated for temperature-dependent processes of the life cycle of *G. p. gambiensis* combining published and new observed data (Fig. S4). The log-linear function for adult mortality (Table S2) differed from published ones of other species (Fig. S4a). Up to 24°C, female mortality rate was 0.013 day^−1^, and mortality increased exponentially with increasing temperatures to reach 0.023 day^−1^ at 32°C, which corresponded to a lifespan of 43-77 days. Male mortality was higher than female mortality (Table S2, Fig. S5).

Pupa emergence clearly followed a logistic function when fitted to the observed data, providing a different pattern as compared to Hargrove’s equation (2004) (Fig S4b, Eq. 14, Table S2).

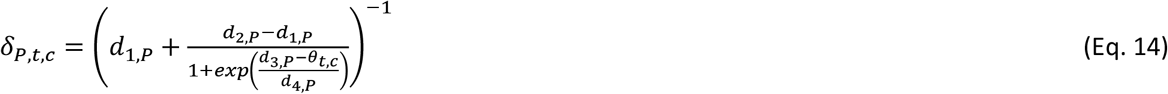

Mark-release-recapture data indicated a dispersal range *r* of one cell, the daily average distance covered proved to be less than 250 m (Fig. S3).

Finally, the spatial heterogeneity of carrying capacities was high, the local density ranging from 112 to 104,768 flies per km² (median: 2,320). On the contrary, spatial variations of local temperatures were small, the standard deviation over the simulated landscape never exceeding 0.67°C at any time step.

### No-control scenario

The no-control scenario was closely in line with field observations made before the start of the Niayes’ control program (Fig. 2). Population dynamics were seasonally influenced (Fig. 2B) and driven by temperature as expected (Fig. 2A). The female fly population dynamics (T+F_1:4+_) was similar across years (Fig. 2B) with a growth rate of −0.75% during the last simulation year. On average 40% of the parous young females (1, 2 or 3 ovulations) had deposited one larva, whereas 33 and 26% of the females had deposited 2 and 3 larvae, respectively (Fig. 2C-D). The spatial variability of age structure (not shown) was 3 to 4 times lower than its temporal variability.

**Figure 2.**
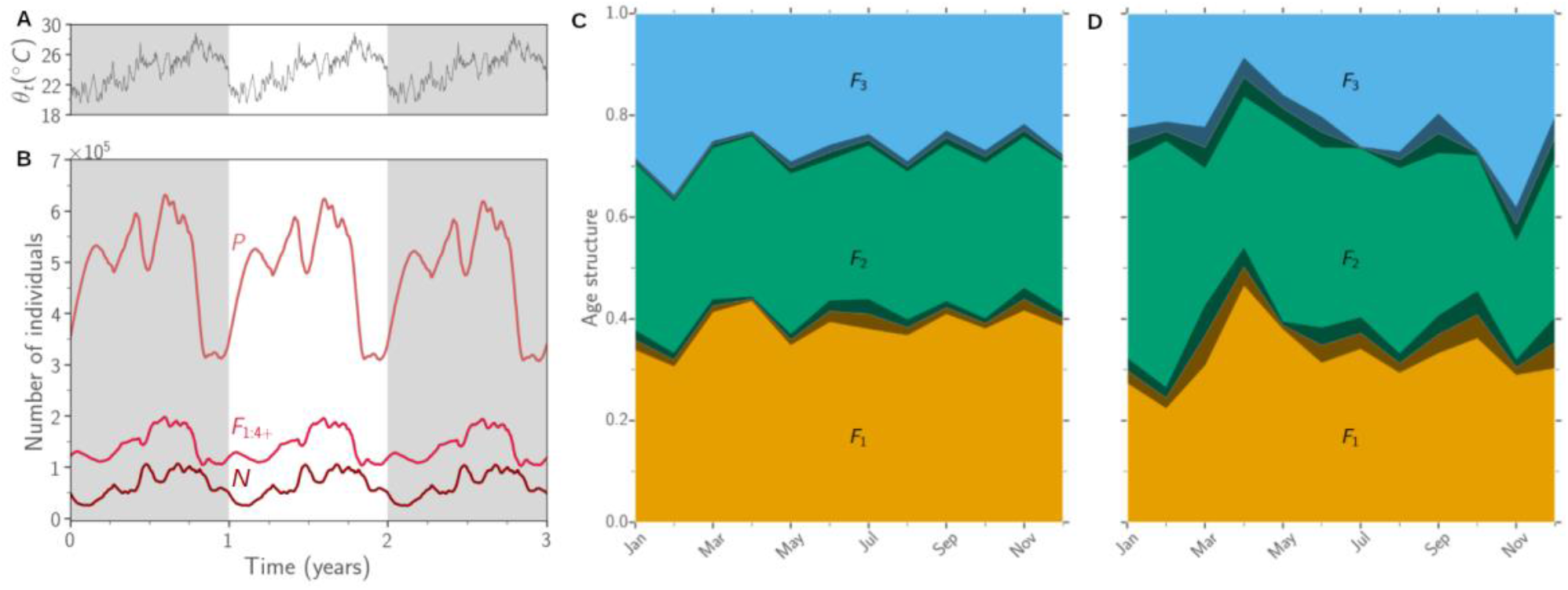
Model predictions for the no-control scenario. A: average daily temperatures over three years (in °C); B: total number of individuals per stage (P: pupae, N: nulliparous females, F: parous females) in the grid (56.25 km^2^) over three years of simulation; C: female age structure 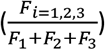 during the last year of simulation; D: observed female age structure (captures and dissection occurred from 2008 to 2011 in the Niayes; results were averaged by month, all years and locations aggregated; grey filled areas are confidence intervals around the mean: 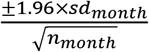, with *sd*_*month*_ the standard deviation and *n*_*month*_ the number of measures, i.e. the number of days in the month for simulations, the number of captures for data).

### Temperature and mortality as key factors driving population size

Model predictions other than age structure (Fig. S7) were highly sensitive to variations in temperature (*θ*) and adult mortality (*μ*_*{N,F,M}*_), and moderately to variations in nulliparous (*δ*_*N*_) and parous (*δ*_*F*_) female larval development duration (Table S4), while parameters related to pupae (*μ*_*P*_, *δ*_*P*_), carrying capacities (*k*), and dispersal (*g*) did not contribute to output variance (Fig. 3, Fig. S8). A 5% variation in temperature resulted in demographic explosion or extinction, substantially outweighing the effect of a similar variation in carrying capacity (Fig. 3A), reinforcing the need for considering reasonable temperature variations. Temperature and adult mortality explained 78% of population size variance (Fig. 3B). Development of nulliparous and parous females added up to another 14.5% of explained variance. Unexpectedly, interactions between parameters were not important.

**Figure 3.**
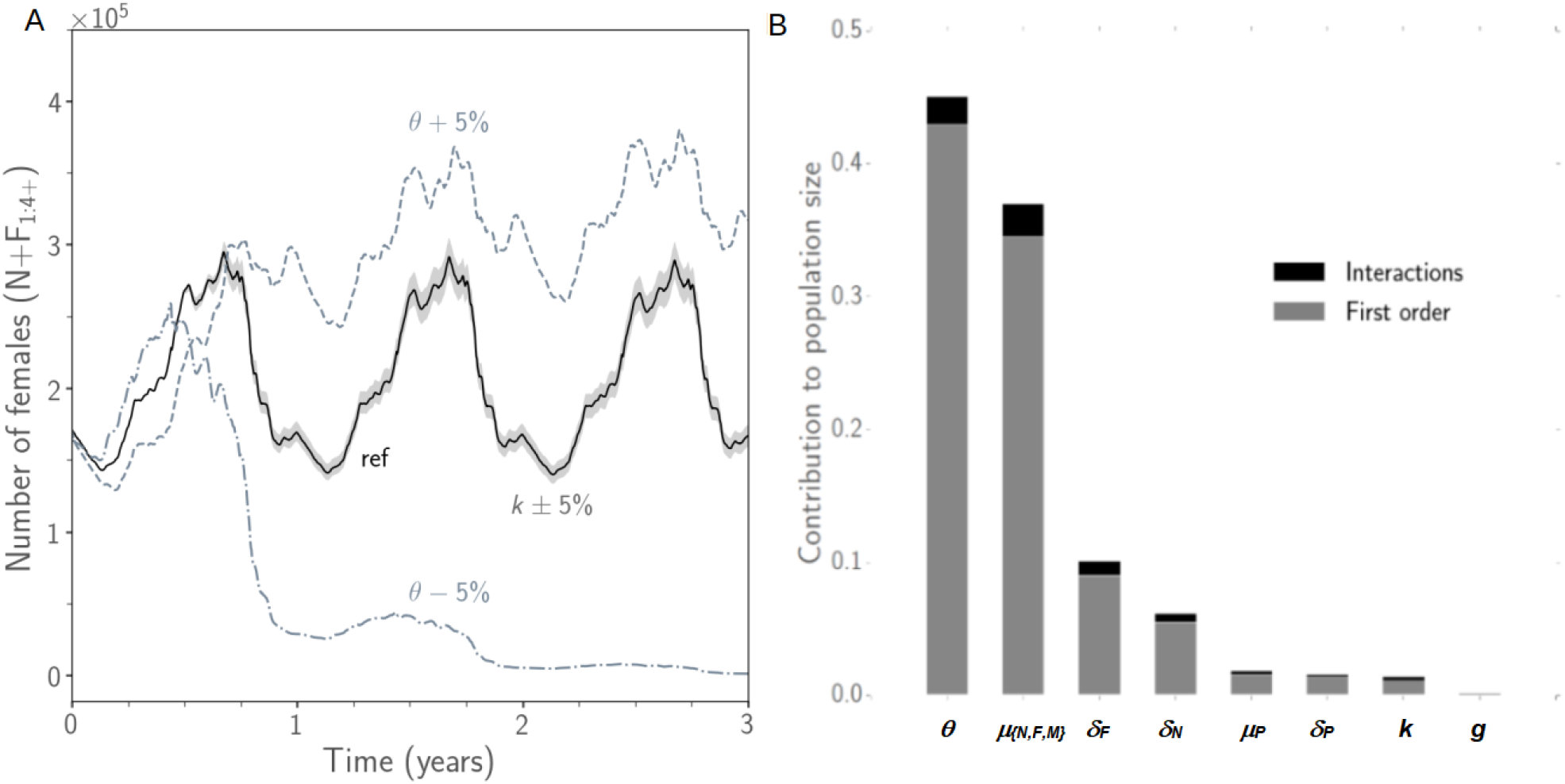
Sensitivity analysis of the model. A: effect on population size (nulliparous and parous females) of temperature variations (+5% from reference: dashed, −5%: dash dot) compared to ±5% variations in carrying capacities (grey filling); B: results of the FAST sensitivity analysis with contribution to population size variance of model parameters (*θ*: temperature, *μ*_*{N,F,M}*_: adult mortality, *δ*_*X*_: time to development of stage *X* (with X in {*P*: pupae, *N*: nulliparous females, *F*: parous females}), *k*: carrying capacity, *g*: the shape parameter in the diffusion process; sensitivity indices for principal effect in grey and for interactions in black). All parameters were varied by ±5% from their reference value except temperature varying by ±0.3°C.

### Efficacy of control measures driven by environmental heterogeneity

Increasing adult mortality to levels comparable to those obtained during control programs (Hargrove 2003) induced a sharp decline in the tsetse population after one year of control (Fig. 4). To obtain a reduction in population size in the simulated area down to 2% of its size without control while applying a homogeneous increase in mortality over space (orange point labelled “2” in Fig. 4A), female life expectancy had to be reduced from 60 (no control) to 35 days. The same cells contributed the most to the total population size irrespective whether control was implemented (Fig. 4B2) or not (Fig. 4B1), and this was closely correlated to local carrying capacity. Upon reaching a low average population density over the area (58 female flies per km^2^), new patterns emerged related to cell-specific properties. Surprisingly, a homogeneous increase in adult mortality had a heterogeneous impact at the cell level: the decrease in local relative population density (i.e. the local control efficacy, Fig. 4C2) was not correlated with the carrying capacity (Fig. S9A), but was correlated with local temperature, i.e. the coldest cells that experienced the smallest variations in temperature showed the lowest impact (Fig. S9B-D). This pattern was obvious despite the small variations in mean temperature (23.7°C to 24.3°C) and standard deviation (1.98°C to 2.37°C). These two temperature statistics were not correlated (Fig. S9C-D).

**Figure 4.**
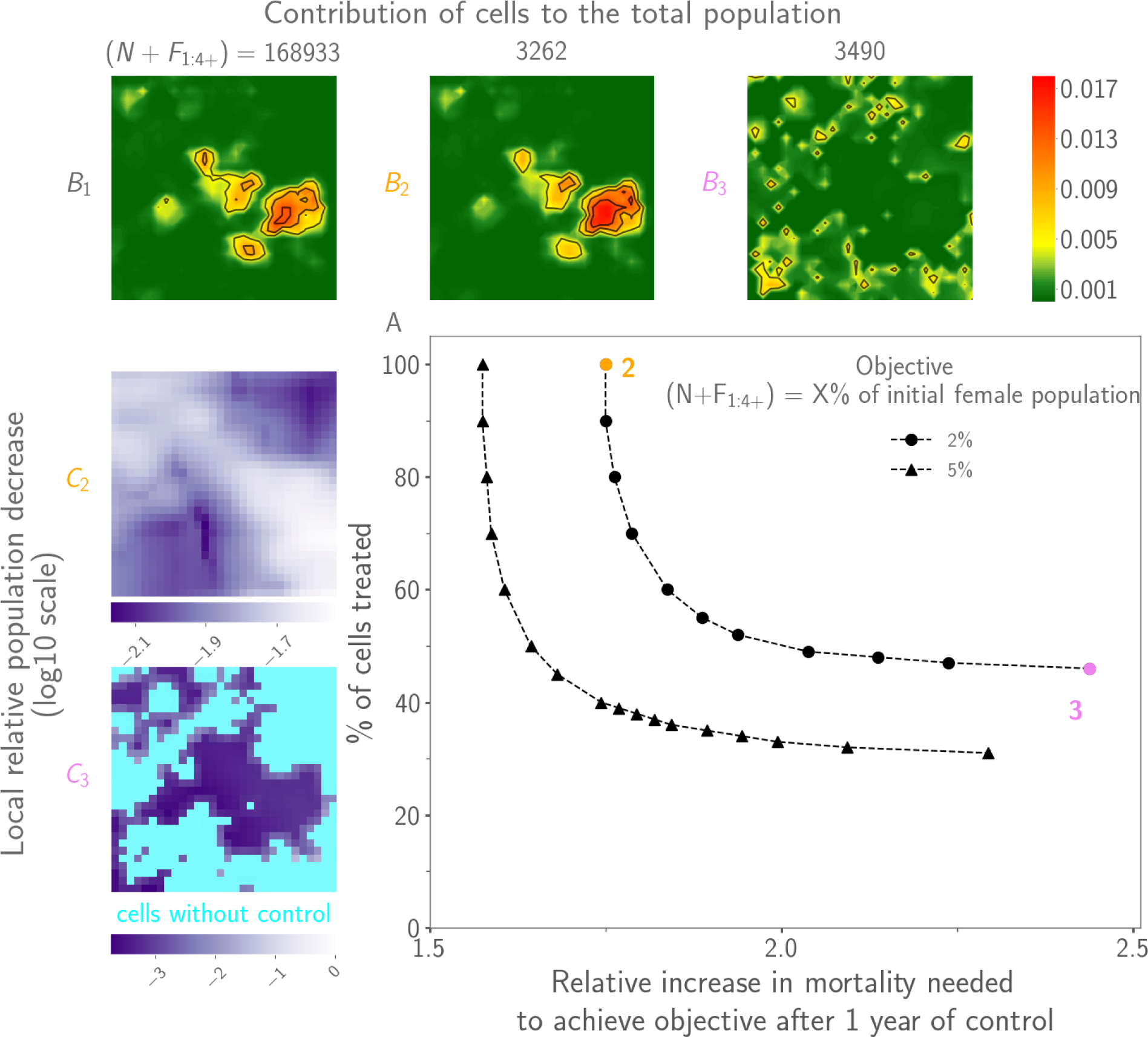
Impact of increasing adult mortality on tsetse fly population size. A: relative increase in mortality needed to achieve a reduction of the female population size to 2% (circle) or 5% (triangle) of its initial size after one year of control, when a fraction of cells was targeted. B: contribution of cells to the total population size (1: no control, 2: homogeneous control, 3: heterogeneous control targeting 46% of the cells). C: local control efficacy (2-3: same as in B), the darkest being the most effective (in cyan, cells without control).

In contrast, targeting cells contributing the most to population management, i.e. those with the greatest carrying capacity and which were most impacted by an increase in mortality, resulted in a similar decrease in population size as a homogeneous control, while requiring a reasonable increase in mortality. However, it resulted in a much more fragmented population and control efficacy was no longer related with temperature. Controlling 70% of the area was as efficient as controlling the whole area (Fig. 4A). Reducing further the proportion of controlled cells required a sharp increase in mortality to obtain a similar efficacy. To obtain a reduction in population size in the simulated area to 2% of its original size without control while applying a heterogeneous increase in mortality (pink point labelled “3” in Fig. 4A), female life expectancy had to be reduced from 60 (no control) to 25 days in 46% of the simulated area (i.e. the average life expectancy over the area was 44 days, slightly higher than in the homogeneous case). If less than 46% of the surface was controlled, the population could not be decreased below 2% of its initial size. Cells contributing the most to the total population size were scattered in the area (Fig. 4B3). The local relative population decrease (Fig. 4C3) here was slightly associated with carrying capacity (Fig. S10A) and the control was more effective in cells with intermediate carrying capacity than in cells with lower ones. However, no effect of temperature was observed (Fig. S10B-C). Similar patterns were observed if the population was to be reduced to 5% of its initial size (not shown).

### Population resurgence after control

Population resurgence one year after the control period was slow irrespective of whether the control was homogeneous or heterogeneous, but resurgence could be high locally in refuges.

After a homogeneous control, a yearly rate of population growth of 23% was observed at the grid scale (from 3,262 to 4,011 individuals). The speed of the resurgence was spatially heterogeneous (Fig. 5A) and growth rates were highest and positive in refuge cells (i.e. coldest cells with lowest temperature variations, Fig. 5C-D), where the impact of control effort was previously the lowest (brown symbols). One year after control, local growth rates were still negative in cells where the control had been the most effective (green and blue symbols). Carrying capacity did not impact resurgence (Fig. 5B).

**Figure 5.**
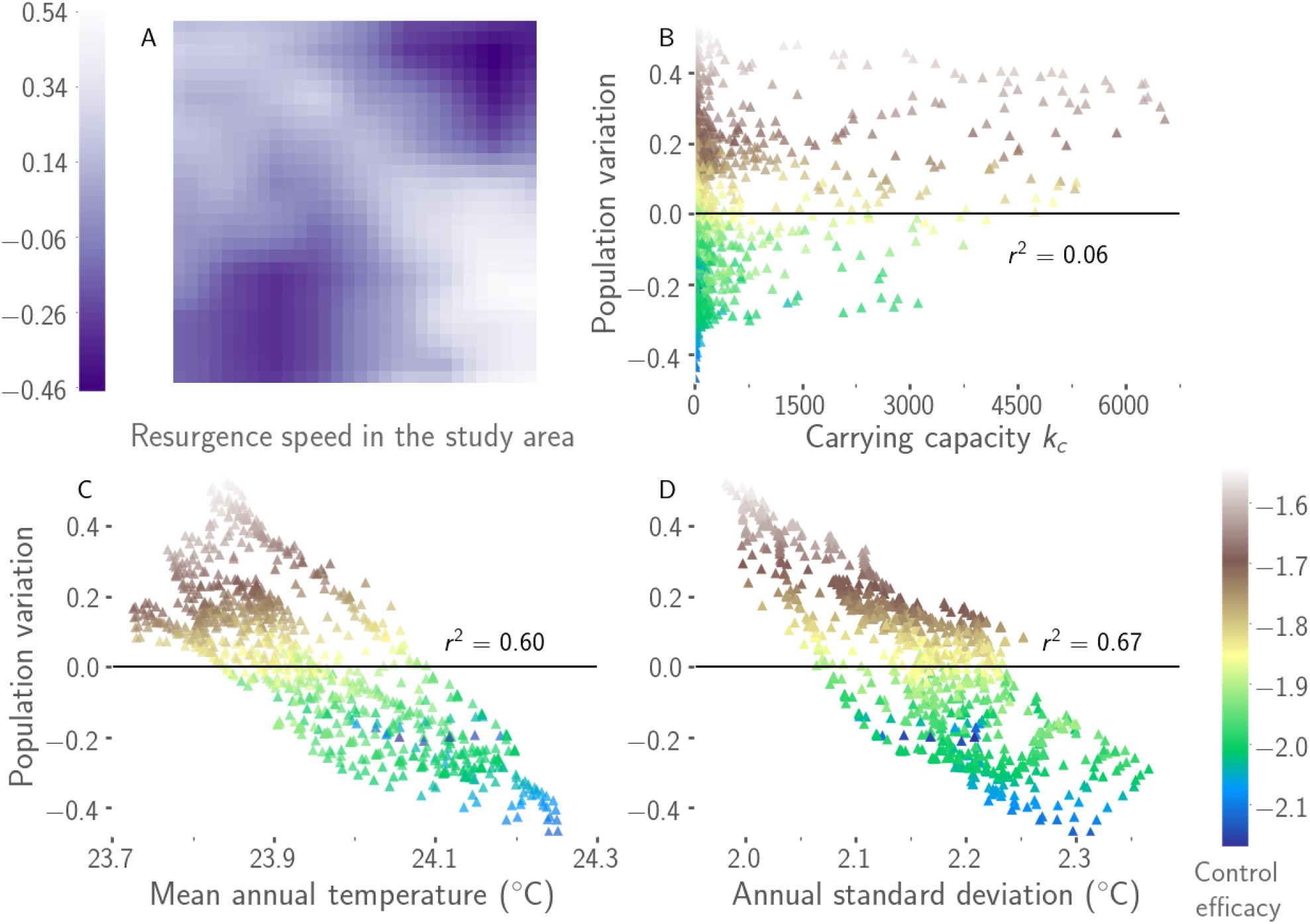
Local population resurgence one year after the end of a one-year homogeneous control. A: local growth rate in the study area. B-D: variations of the local growth rate with carrying capacity, the mean annual temperature, and the annual standard deviation of temperature. Colours in B-D represent control efficacy with blue being the most effective.

After a heterogeneous control, the yearly population growth rate was lower as compared with a homogeneous control effort, with only a 1% population increase at the grid scale (from 3,490 to 3,528 individuals). Unexpectedly, such a control resulted in contrasting situations with very high local growth rates in a few cells (Fig. 6A), without any correlation with local characteristics or with the scores used to target controlled cells (Fig. 6B-D). Refuges were located at the interface between controlled and uncontrolled zones (Fig. 6A), and monitoring efforts after the control period should particularly focus on cells of intermediate carrying capacity (Fig. 6B).

**Figure 6.**
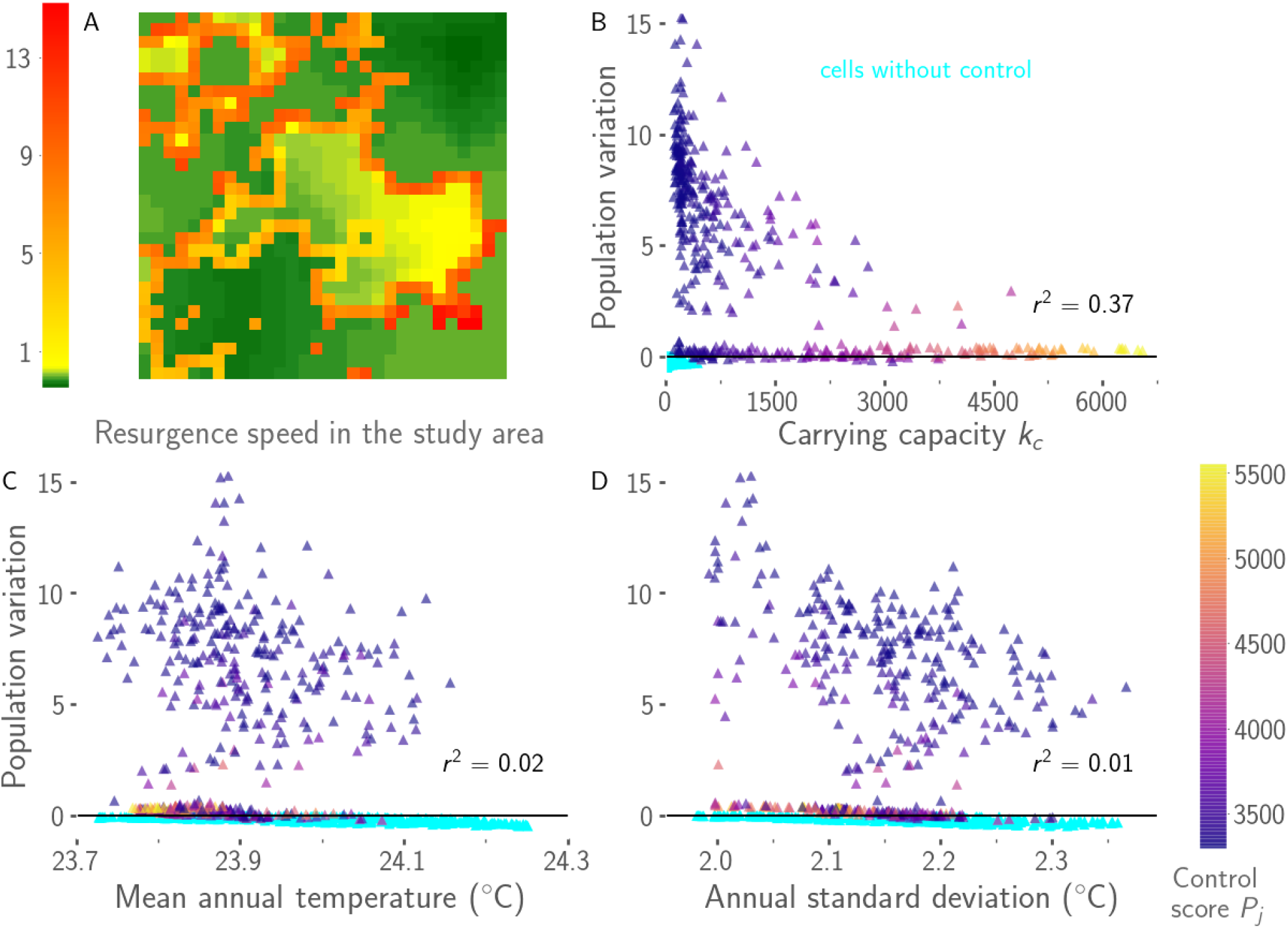
Local population resurgence one year after the end of a one-year heterogeneous control (46% of controlled cells). A: local growth rate in the study area. B-D: variations of the local growth rate with carrying capacity, the mean annual temperature, and the annual standard deviation of temperature. Colour in B-D represents control score, the highest being still targeted when the proportion of controlled cells is reduced. In cyan: uncontrolled cells (have lower scores).

## Discussion

Environmental heterogeneity with respect to carrying capacity and temperature not only drove the temporal population dynamics of *G. p. gambiensis* at large scale, but also the spatial distribution of individuals, as well as control efficacy. It unexpectedly rendered heterogeneous the impact of a homogeneous increase in adult mortality on population dynamics. The coldest cells with the smallest variations in temperature acted as refuges when adult mortality was homogeneously increased, and in these refuges, the control effort was less effective and population resurgence faster after control had stopped. Such a heterogeneous impact can be partially compensated during eradication campaigns by releasing sterile males by air that will aggregate in the same sites as wild males, as observed in the eradication campaign against *Glossina austeni* on Unguja Island of Zanzibar (Vreysen et al. 2011). To increase the chances of success, control strategies should account for environmental heterogeneity and emphasise (1) local areas of high suitability characterized by a high carrying capacity, (2) local refuges characterized by lower local temperatures within the relevant range for tsetse (23.7-24.0°C here), and (3) local areas with low variability of temperature over the year (irrespective of carrying capacity). In contrast, targeting patches where population control is the most efficient enabled to decrease population size with a similar efficacy, but this approach resulted in much more dispersed individuals, and in addition, efficacy of the control effort was no longer related to temperature. In this case, population resurgence after control, while being very slow in general, was locally very high in refuges, which differed from previous refuges in that they were located on the interface between controlled and uncontrolled zones. Refuges, highlighted in our study area despite a small surface suitable for tsetse, could jeopardize control efforts by providing areas from which recolonization may occur after control has stopped, a result that was at the origin of the principle of area-wide pest management by Knipling (Vreysen et al. 2011).

In addition, the temperature effect on tsetse population dynamics both at a large and small local scales emphasises the need for further investigating the impact of climate change on tsetse populations (Terblanche et al. 2008; Moore et al. 2012). It is unlikely that tsetse flies will cross the Sahara, but they could migrate to higher altitudes and invade trypanosoma-free zones, particularly in Eastern and Southern Africa where tsetse distribution is mainly governed by altitude (Solano et al. 2010a). Such population shifts will impact the density of cattle in either direction by impacting the transmission of trypanosomoses, which may in turn impact the distribution of wild fauna including lions (Carter et al. 2018). In addition, isolated populations could merge if close enough together in a changing habitat, possibly impairing control strategies. Conversely, new populations could become isolated, all the more so as temperature is the first driver of landscape friction in tsetse (Bouyer et al. 2015).

The mechanistic spatio-temporal model developed to predict *G. p. gambiensis* population dynamics and how these evolve when adult mortality is increased is original compared to already published models. First, the model incorporated environmental heterogeneity through a data-driven approach, both accounting for variable temperatures and carrying capacities in space and time. The model used realistic assumptions and highlighted the importance of refuges in this species, which was not previously evidenced using theoretical assumptions (Childs 2011), knowledge-driven patterns (Barclay & Vreysen 2013), or aggregated patterns assuming a binary occupancy (Lin et al. 2015). The proposed model can be applied to other areas with available data and a known metapopulation structure. Second, recent field and laboratory data on mortality, development, and dispersal were incorporated into the model. Predicted age structure was in good agreement with field data, and proved robust in our simulations as it was barely impacted by parameter variations. Amplitude and duration of seasons are expected to be major drivers of ovarian age distribution, but this could not be assessed here as temperature data were only available for one year. Our results highlighted the need for more biological studies to better infer mortality variations with temperature, as well as the need for innovative methods to more accurately estimate temperatures as perceived by the insects. Such a complementary interplay between models, field observations, and laboratory experiments is fundamental to make accurate predictions.

The fact that tsetse fly population dynamics was much more sensitive to mortality than reproduction is consistent with tsetse flies being specialists with a narrow niche. In this species, individual survival is prioritized over breeding (Pagabeleguem et al. 2016), where other species compensate for losses by boosting birth rates (Southwood et al. 1974). *Glossina* spp. have evolved towards an optimal utilization of energy and resources (Cody 1966), which makes them highly adapted to their ecological niche. Therefore, they are less likely to leave their habitat and expose themselves to other environments, which keeps the population at or near carrying capacity (Southwood et al. 1974).

Efficient control methods have to be designed considering the ecological strategy of the concerned species (Southwood et al. 1974; Conway 1977). Fast action methods such as chemicals are better suited for species showing high reproductive rates, short generation times, along with broad food preferences and good dispersing abilities (Altieri et al. 1983). In contrast, pests reproducing at lower rates and having longer generation time but good competitive abilities would be more efficiently restrained with cultural control, host resistance, and sterilization (Altieri et al. 1983). Nonetheless, such quite extreme characteristics should be considered in conjunction with species relationships within communities (Ehler & Miller 1978; Altieri et al. 1983).

Traps, targets, and insecticide-treated livestock are control tactics increasing adult mortality, which can drastically reduce tsetse populations (Kagbadouno et al. 2011; Dicko et al. 2014; Percoma et al. 2018). However, our results indicate also generation time as a contributing factor to population size variations. Such a factor can be indirectly modified using the sterile insect technique, which impair reproduction (Dyck et al. 2005). Obtaining very low tsetse densities is not enough to reach eradication as was demonstrated recently against *G. p. gambiensis* in north-western Ghana (Adam et al. 2013), the Loos islands in Guinea (Kagbadouno et al. 2011), and the Mouhoun river in Burkina Faso (Percoma et al. 2018). In addition, in view of unexpected local refuges where increasing adult mortality is not as effective as in other areas, it becomes necessary to further assess the effect of combined and spatially targeted control measures to achieve eradication.

Our model provides a relevant tool to evaluate complex control strategies as it accounts simultaneously for density-dependent processes, spatial and temporal environmental heterogeneity, and all stages of tsetse lifecycle possibly targeted by control measures. Our framework could also be useful to identify where to focus stakeholders’ efforts to minimize impact of other specialist pests, such as the codling moth (*Cydia pomonella*) affecting apple and pear trees, and the sheep ked (*Melophagus ovinus*). Nevertheless, the importance of stochastic events when populations become very small must not be overlooked and these effects should be included in future developments. Our approach gives clues on how to trigger a drastic decline of the population. However, to predict the subsequent population dynamics at low densities and assess final steps of eradication strategies, a deterministic framework becomes irrelevant as it does not enable quantifying the probability of population extinction at local and large scales.

Accounting for spatial heterogeneity is essential to better understand and predict tsetse population dynamics, as habitat fragmentation holds the key to population survival when conditions are globally hostile. However, parameters driving tsetse fly dispersal abilities did not structure their final distribution. Landscape ecology must be studied to identify patches that will need longitudinal surveillance. Optimal management strategies are therefore valid for a given species in a given habitat and should not be generalized without baseline data collection to characterize the ecosystem.

To conclude, environmental carrying capacity largely explained the contribution of local source spots to tsetse fly population dynamics at a large scale, but unfavourable conditions resulted in a progressive disappearance of such spots and the existence of refuges that located in colder areas where the temperature was less variable. When applying a spatially homogeneous increase in adult mortality for one year, population size was less impacted in such refuges. In contrast, applying a spatially heterogeneous increase in adult mortality resulted in refuges located at the interface between controlled and uncontrolled zones, and previous temperature-dependent refuges disappeared. Areas to be controlled should be chosen with caution when facing a heterogeneous habitat. Our study confirmed the importance of a preliminary characterization of the study area before the start of control operations in order to include the most suitable habitats in the control strategy, which is the foundation of area-wide integrated pest management.

## Data accessibility

Data are available in the Supporting Information and online: https://sourcesup.renater.fr/projects/spatial-tsetse/

## Supplementary material

Script and codes are available online: https://sourcesup.renater.fr/projects/spatial-tsetse/

## Authors’ contribution

JB and PE designed the study and advised biological details. HC, SA, SPi and PE developed the model. HC conducted the analyses and prepared the figures. HC, SA, SPi, JB and PE discussed the results. HC and PE wrote the manuscript. AD provided model external input data readily usable by the mechanistic model. JB, MTS, BS, MB, MV, SPa, AB collected the data. All authors edited the manuscript.

## Acknowledgements

This preprint has been peer-reviewed and recommended by Peer Community In Ecology (https://dx.doi.org/10.24072/pci.ecology.100024). The authors are thankful to the technicians of the vet services from Senegal and ISRA for collecting the field data used in this study.

## Appendix

[This is the link to Supporting Information](https://www.biorxiv.org/content/10.1101/493650v2.supplementary-material).

