## Supporting Information for "Environmental heterogeneity drives tsetse fly population dynamics and control"

#### 1. Experimental data

**Table S1.** Pupa emergence. One hundred and twenty 20-day old pupae were hold in climate controlled rooms until emergence. The experiment was replicated three times for each temperature tested.

| | Phase 1 | | | Phase 2 | | | |
|---|---|---|---|---|---|---|---|
| | Temperature (°C) | Duration (days) | Daily development rate ($day^{-1}$) $.10^{-2}$ | Temperature (°C) | Time before hatching (days) | Number of flies | Daily development rate ($day^{-1}$) $.10^{-2}$ |
| Females | 25 | 20 | 3.69 | 15.00 | 24.37 | 63 | 1.08 |
| | | | | 15.00 | 22.53 | 57 | 1.16 |
| | | | | 15.00 | 22.24 | 66 | 1.18 |
| | | | | 20.11 | 19.19 | 54 | 1.37 |
| | | | | 20.11 | 18.95 | 60 | 1.39 |
| | | | | 20.11 | 18.80 | 64 | 1.40 |
| | | | | 25.40 | 6.13 | 48 | 4.29 |
| | | | | 25.41 | 7.89 | 45 | 3.33 |
| | | | | 25.80 | 6.46 | 61 | 4.06 |
| | | | | 27.50 | 5.80 | 66 | 4.52 |
| | | | | 27.50 | 5.72 | 65 | 4.59 |
| | | | | 27.50 | 5.55 | 58 | 4.73 |
| Males | 25 | 20 | 3.45 | 15.00 | 27.91 | 43 | 1.11 |
| | | | | 15.00 | 27.12 | 49 | 1.15 |
| | | | | 15.00 | 26.96 | 53 | 1.15 |
| | | | | 20.11 | 23.19 | 58 | 1.34 |
| | | | | 20.11 | 22.35 | 55 | 1.39 |
| | | | | 20.11 | 22.15 | 52 | 1.40 |
| | | | | 25.40 | 8.20 | 66 | 3.79 |
| | | | | 25.41 | 10.70 | 66 | 2.90 |
| | | | | 25.80 | 8.60 | 58 | 3.61 |
| | | | | 27.50 | 7.13 | 48 | 4.36 |
| | | | | 27.50 | 7.00 | 52 | 4.44 |
| | | | | 27.50 | 7.00 | 55 | 4.44 |

#### 2. Additional support to model input data

##### 2.1 Carrying capacities

Suitability Index (SI) - The first layer needed to estimate the carrying capacity is the habitat suitability index. We used this layer to determine the area where tsetse flies can survive (suitable habitats). A statistical analysis of the habitat was carried out using correlative species distribution models. The methodology used is based on the framework developed in the Niayes (Senegal) using the Maximum Entropy model (MaxEnt) (Dicko et al. 2014). MaxEnt is one of the most widely-used species distribution models. It is a machine learning method based on the information theory concept of maximum entropy (Elith et al. 2011). It fits a species distribution by contrasting the environmental condition where the species is present to the global environment characterized by some generated pseudo-absence data, also called the background. Occurrence data from already described entomological surveys were used as input. Characterization of the environment in the study area relied on the 5-year average, minimum, maximum, range, and standard deviation of four spatio-temporal layers (day land surface temperature (DLST), night land surface temperature (NLST), normalized difference vegetation index (NDVI), maximum middle infrared (MIR)), with the digital elevation model (DEM) added to the set of summarized variables. To account for the sampling bias present in the entomological data, a gaussian kernel based grid that gives more weight to more densely sampled areas was needed (Bouyer et al. 2009). Model complexity in the MaxEnt framework can be controlled using the beta regularization parameter. Five parameters (1, 1.5, 2, 3, 4) were tested. We performed multimodel inference using model averaging weighted by the Akaike Information Criterion (AIC) to choose the best model (Burnham & Anderson 2002; Warren & Seifert 2011). The final output was a suitability index that ranged between 0 (least suitable habitat) and 1 (most suitable habitat) defined for every patch, providing a quantitative indicator of the habitat preferences of *G. p. gambiensis* in the study area.

Apparent Density per Trap (ADT) - The second layer needed to estimate the carrying capacity is the apparent density of tsetse flies per trap per day (ADT), as measured using biconical traps considered here as substitution hosts (Dicko et al. 2015). ADT is considered as an apparent density because it does not depend only on the real fly density but also on fly dispersal, feeding frequency, and age structure which all impact trap catches. We predicted a 5-year average of tsetse ADT at a spatial resolution of $1km^2$ using a geostatistical model fitted against the computed suitability index. A negative binomial model with spatial random effects was used. Negative binomial models can be seen as an extension of the classical Poisson regression to account for over-dispersion in count data. In addition, because of the sampling bias and the clustering of observations in such entomological dataset, a spatial random effect using the Matern correlation structure was used (Cressie & Cassie 1993).

##### 2.2 Temperature

We downloaded high spatial resolution (1km) daily air temperature data in the region from the Meteo1Km project (http://dailymeteo.org/content/about). This dataset results from the combination of MODIS Land Surface Temperature and meteorological datasets from around the world through the use of a spatio-temporal geostatistical model (Kilibarda et al. 2014). We then refined these grids to a 250m spatial resolution. However, these temperatures overestimate perceived temperatures and thus will result in erroneous model outcomes. Indeed, meteorological stations are located in areas mostly unsuitable for tsetse flies with higher temperatures than what they experience in their resting places where they stay most of the day. We thus used our own network of temperature data recorded every 15 minutes in some suitable patches and corrected the bias present in the initial data. We ended up with a more accurate model of air temperature in suitable areas that can be used to realistically simulate tsetse fly population dynamics.

### 3. Modelling tsetse fly population dynamics

#### 3.1. Within-cell dynamics

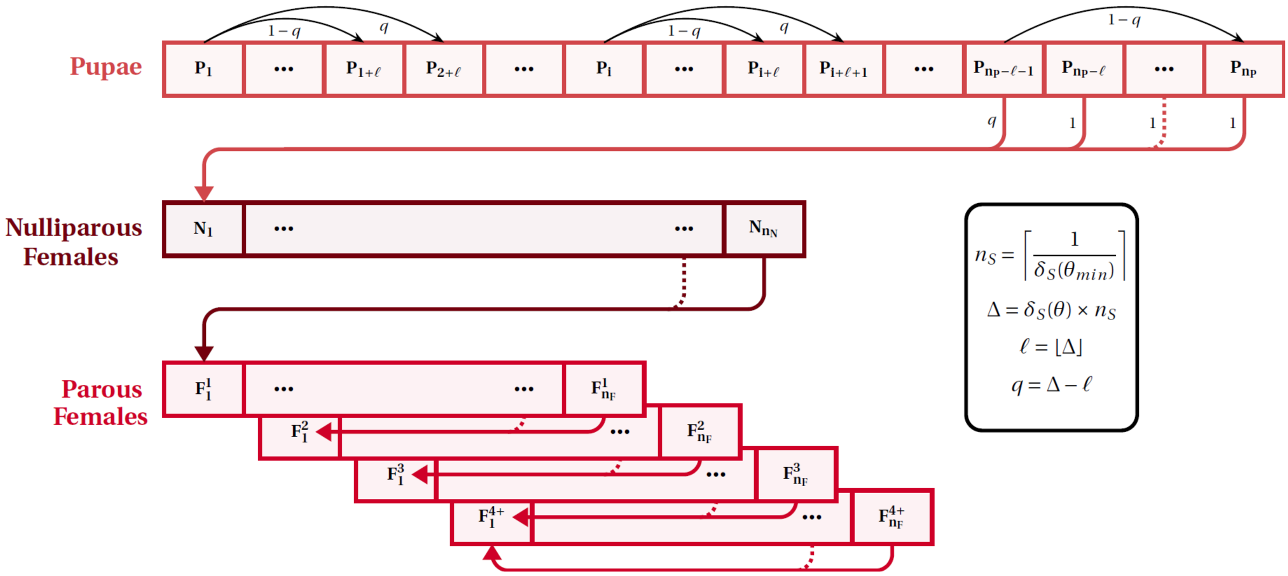

**Fig. S1.** Within-cell model diagram of tsetse fly population dynamics (time unit is a day). All transitions between stages except from pupa (P) to nulliparous female (N) trigger the birth of a new pupa $P_1$. Transitions occur at development rate $\delta_S$ for stage S according to temperature $\theta_{t,c}$ at time $t$ in cell $c$, giving rise daily to a minimum jump of $l$ states from each state i of stage S, with (1-$q$)$S_{t,c,i}$ individuals going from state $S_{t,c,i}$ to state $S_{t+1,c,i+l}$ and $qS_{t,c,i}$ individuals going to $S_{t+1,c,i+l+1}$. If $i + l > n_S$ (respectively $i + l + 1 > n_S$), then concerned individuals go to the next stage. Stage S ∈ {P, N, $F_x$, M}, parity $x$ ∈ {1, 2, 3, 4+}.

#### 3.2. Spatial dispersal

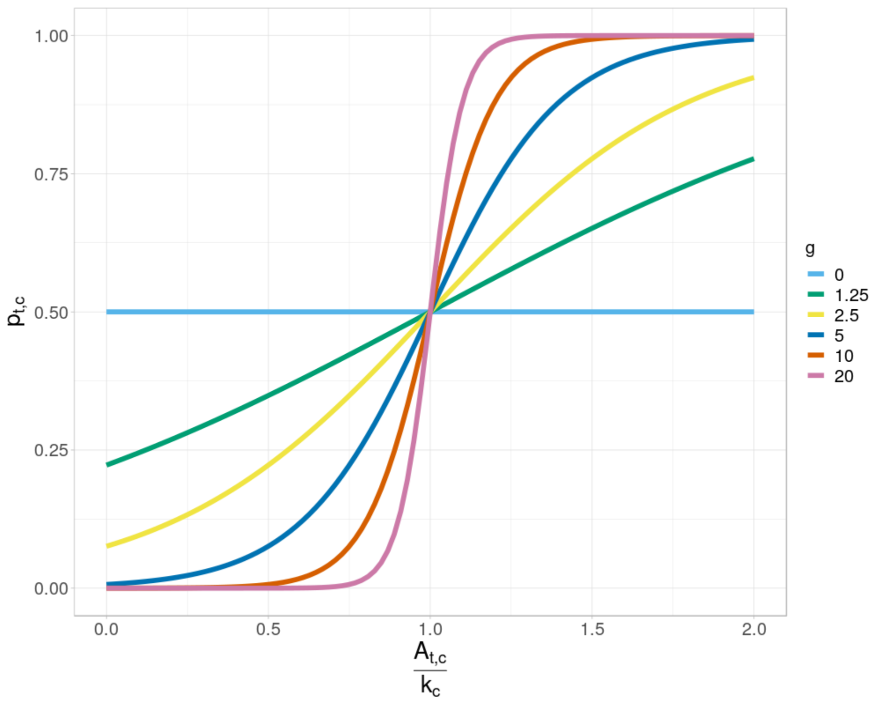

**Fig. S2.** Dispersal rate $p_{t,c}$ as a function of $\frac{A_{t,c}}{k_c}$ and parameter $g$. The orange line is the equation used in main text ($g$=10). Formula given in main text comes from Lloyd-Smith 2010.

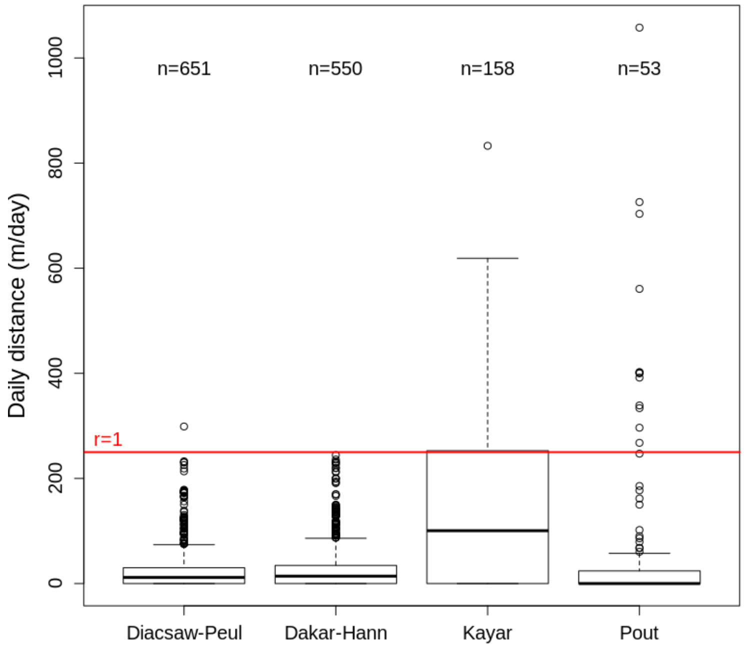

**Fig. S3.** Distance (in meters) between releases and captures of marked tsetse flies, averaged by day for each of the observed locations (Niayes, Senegal).

#### 4. Additional information on model parameters

**Table S2.** Parameter values. These parameters are used in the equations provided in the main text.

| Parameter | Symbol | Value | Standard error / [Reference] |
|---|---|---|---|
| Pupa mortality rate (/day) | $m_P$ | 0.01 | [Childs 2011] |
| Mortality function parameters for parous females | $m_{1,F}$ | 0.358 | 0.004 |
| | $m_{2,F}$ | -12.94 | 0.09 |
| Mortality function parameters for adult males | $m_{1,M}$ | 0.254 | 0.007 |
| | $m_{2,M}$ | -9.53 | 0.19 |
| Development function parameters for pupae | $d_{1,P}$ | 89.0173 | 1.3244 |
| | $d_{2,P}$ | 18.8291 | 2.3717 |
| | $d_{3,P}$ | 22.1831 | 0.2770 |
| | $d_{4,P}$ | 1.7610 | 0.1965 |
| Development function parameters for nulliparous females | $d_{1,N}$ | 0.0020 | 0.0009 |
| | $d_{2,N}$ | 0.061 | 0.002 |
| Development function parameters for parous females | $d_{1,F}$ | 0.0052 | 0.0001 |
| | $d_{2,F}$ | 0.1046 | 0.0004 |
| Shape parameter of the dispersal function | g | 10 | * |
| Cell carrying capacities [min, med, max] | $k_c$ | [6.64 ; 145.65 ; 6547.48] | ° |
| Cell daily temperatures (°C) [min, med, max] | $T_{t,c}$ | [18.66 ; 24.34 ; 29.90] | ° |

*To the best of our knowledge; °Data-driven

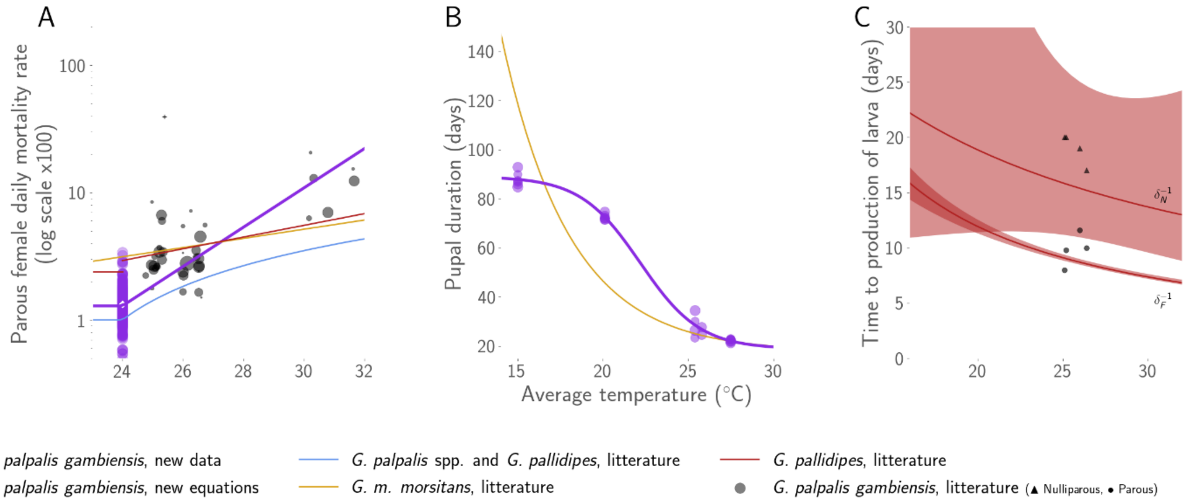

**Fig. S4.** Data (symbols) and predictions (lines) fitted to the new data (if relevant) and from literature for temperature-dependent processes of the model: (a) parous female daily mortality rate (in log-scale); (b) pupal duration (in days); (c) time to larviposition for nulliparous (N, upper curve, triangles) and parous females (F, lower curve, dots). Data from Pagabeleguem et al. (2016) is shown in grey (the cross in (a) was considered an outlier). New data on *G. p. gambiensis* (from FAO/IPCL and CIRDES) is shown in purple, with the barycentre of mortality rate at 24°C highlighted as a white-filled diamond. Purple thick lines are the newly calibrated equations used in the population dynamics model. Predictions from Barclay's equation (2011) is in blue. Orange lines correspond to predictions from Hargrove's equations (2004), with filled areas in (c) corresponding to prediction intervals. Equations for time to larviposition were not modified as only few new data were available, which were consistent with Hargrove's equation.

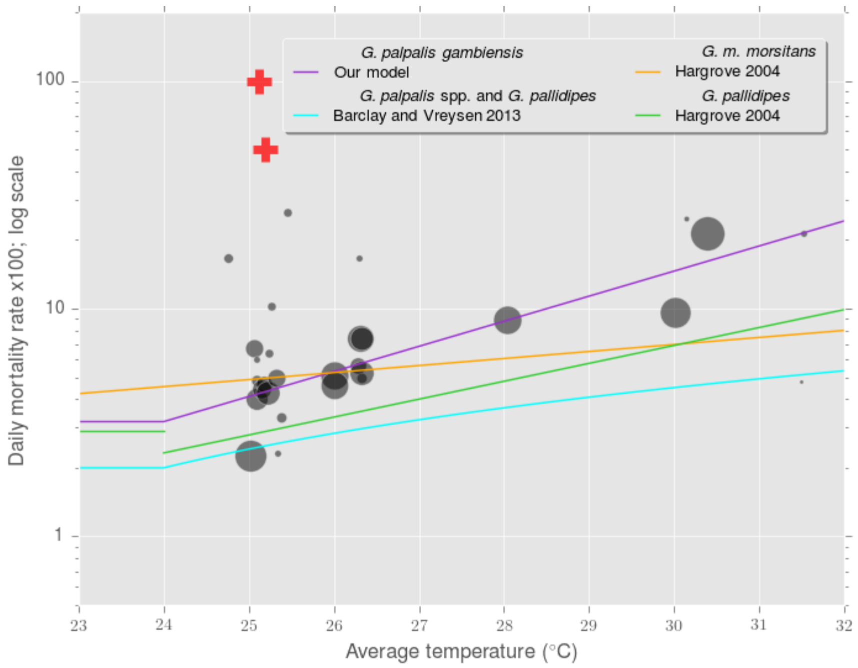

**Fig. S5.** Calibration of male mortality rate. Data and comparison to literature. Red cross: outlier. Point size proportional to sample size.

#### 5. Sensitivity analysis

**Table S3.** Definition of aggregated outputs. All were computed the last year of the simulation.

| Output | Description | Comment |
|---|---|---|
| popMean | Average female population size ($N+F_{1:4+}$) over the year in the grid | Results in main text |
| popStd | Standard deviation of female population size ($N+F_{1:4+}$) over the year in the grid | |
| surfX | Average proportion of cells in the study area with $\frac{\overline{A_c}}{k_c} \geq X$ ($\overline{A_c}$ : average number of adults (nulliparous and parous females, males) over the year in cell $c$) | X = [10,20,50,75,80,90] % |
| surf1 | Average proportion of cells containing at least one adult (nulliparous and parous females, males) on average | |
| miX, meX, maX | Minimum, mean, maximum $\frac{F_x}{F_1+F_2+F_3}$ over the grid and over the year (%) | X = [1,2] |
| mean_distrib | Average $\frac{\overline{A_c}}{k_c}$ over the year | |
| std_distrib | Standard deviation of $\frac{\overline{A_c}}{k_c}$ over the year | |
| qX_distrib | Percentile X of $\frac{\overline{A_c}}{k_c}$ over the year | X = [5,25,75,95] |
| median_distrib | Median of $\frac{\overline{A_c}}{k_c}$ over the year | |

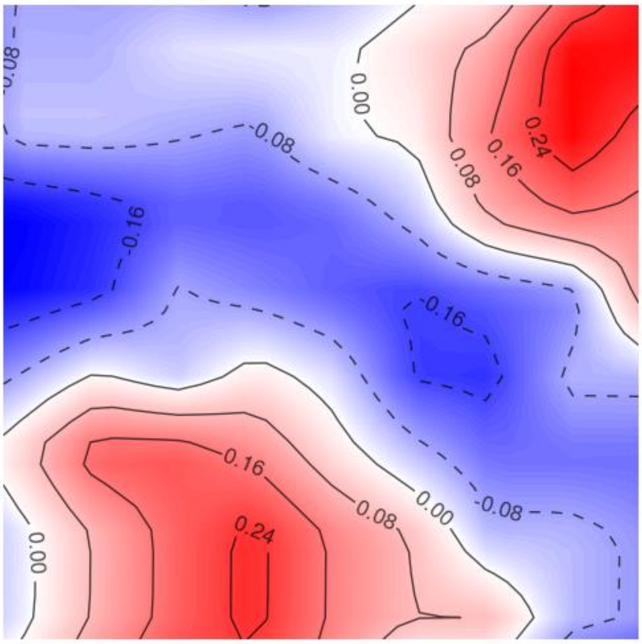

**Fig. S6.** Scaling down of temperature variations for the global sensitivity analysis. Average $T_{cell} - mean_{day}(T) = \frac{\sum_t\left(\theta_{t,c} - \frac{\sum_i \theta_{t,i}}{30*30}\right)}{365}$ over the year for every cell. Maximum is +0.3°C.

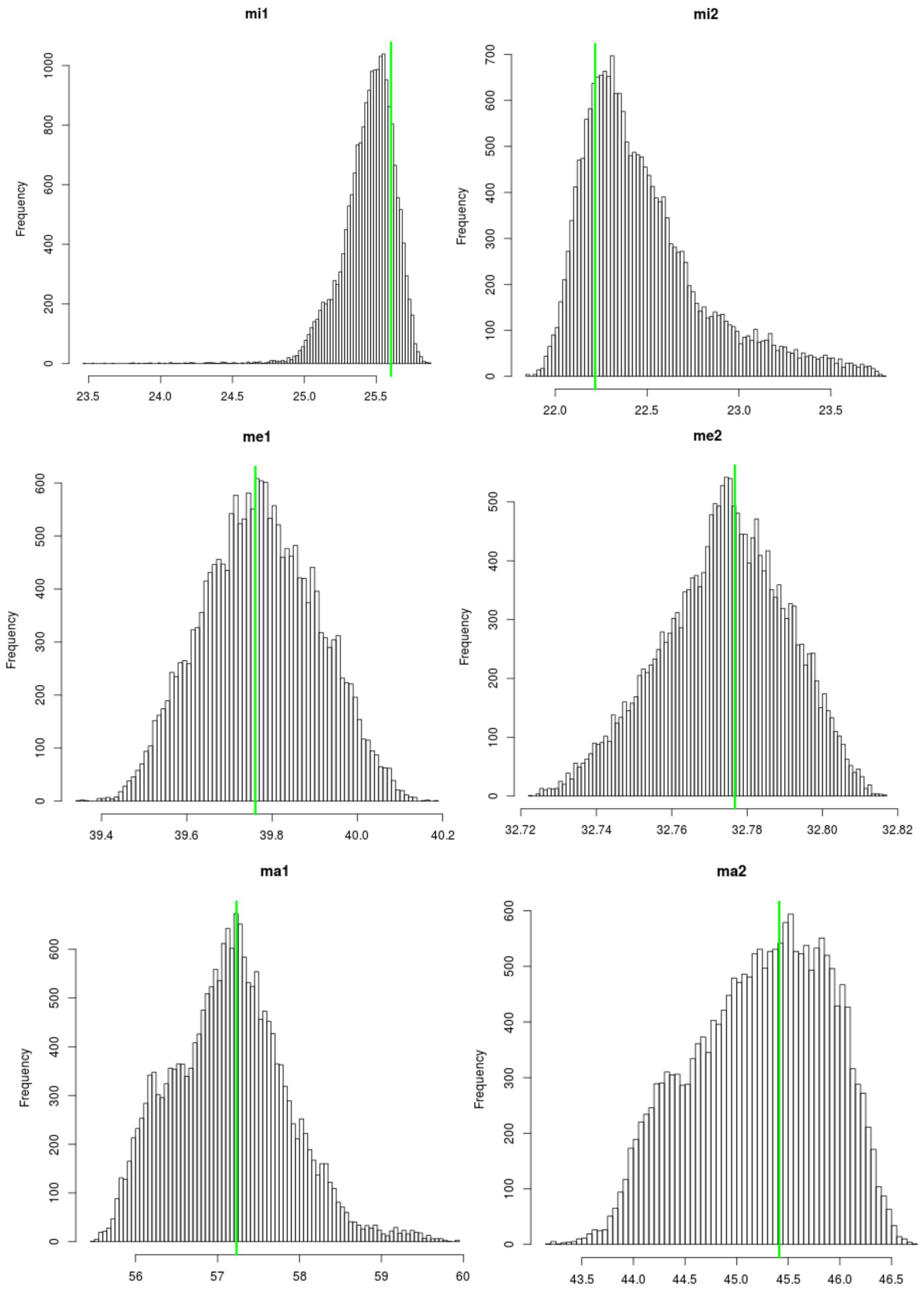

**Fig. S7.** Age structure showed almost no variation with parameter variations, thus was not further analysed. Here, we show variations in the minimum (Mi), mean (Me), and maximum (Ma) proportion of females of parity $X$ ($\frac{F_x}{F_1+F_2+F_3}$, $X$ in [1-2]) over the grid the last year of simulation. Green line: no-control scenario.

**Table S4.** Total sensitivity indices per influential input for outputs showing sufficient variations from their reference value. See definition of outputs Table S3.

| | % of variance explained (sense of variation) | | | |
|---|---|---|---|---|
| Outputs varying significantly | $T$ (-) | $\mu_{\{T,F,M\}}$ (-) | $\delta_F$ (+) | $\delta_T$ (+) |
| popMean | 43.0 | 34.6 | 9.0 | 5.5 |
| popStd | 44.0 | 32.0 | 8.3 | 5.4 |
| surf20 | 35.3 | 37.4 | 10.5 | 5.3 |
| surf1 | 37.8 | 38.6 | 10.3 | 5.1 |
| mean_distrib | 40.5 | 37.6 | 10.3 | 5.8 |
| median_distrib | 38.9 | 35.7 | 9.8 | 5.7 |
| q95_distrib | 41.8 | 32.5 | 9.2 | 5.5 |

popStd

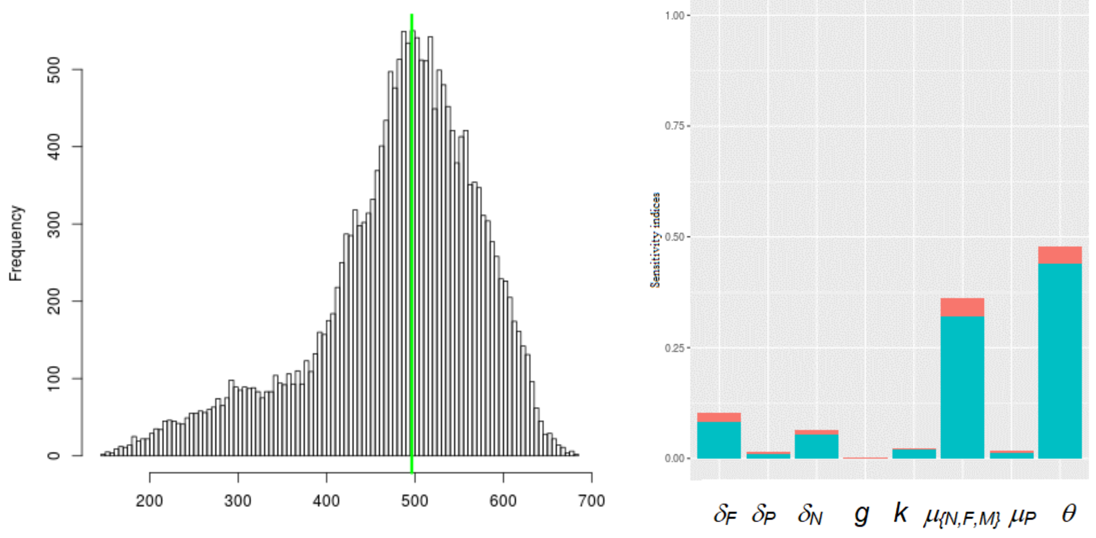

surf20

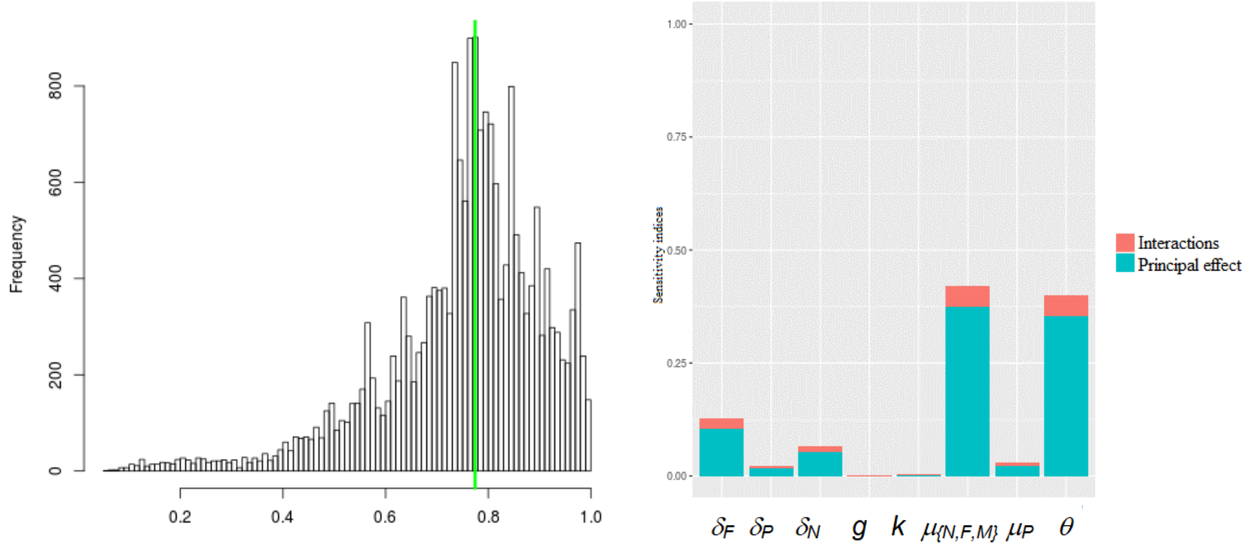

surf1

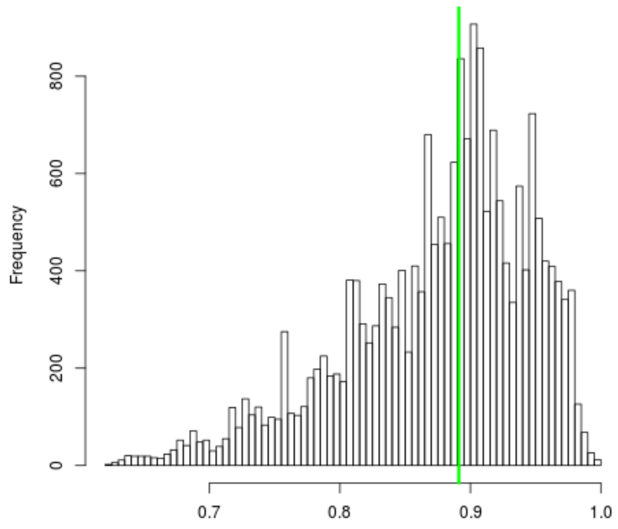

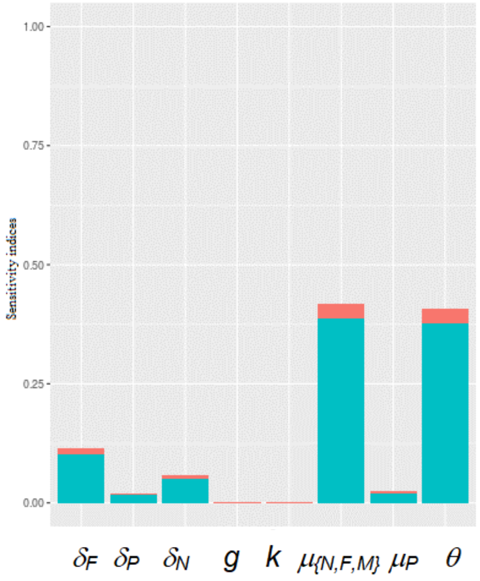

mean_distrib

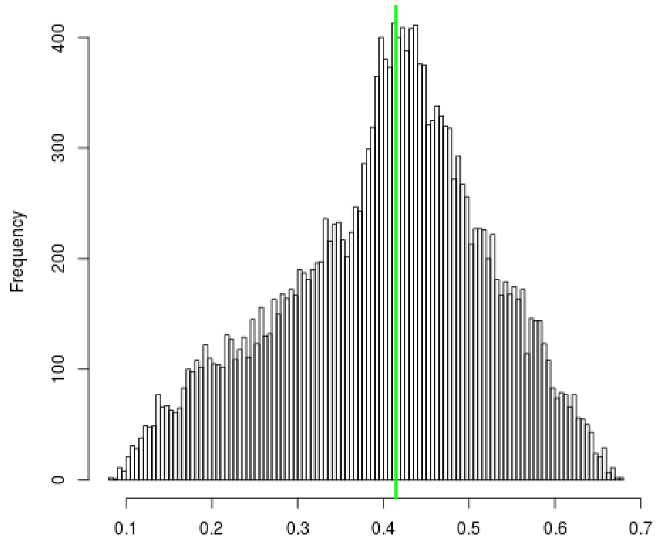

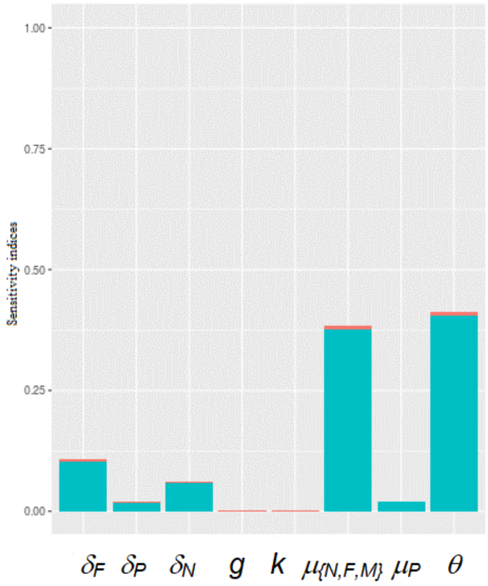

q95_distrib

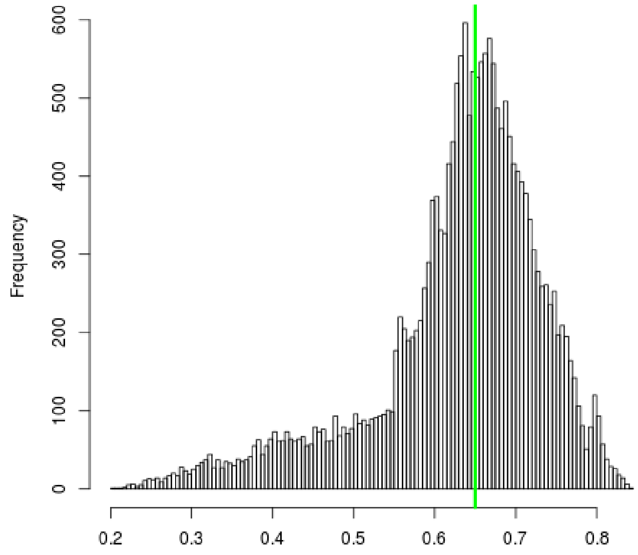

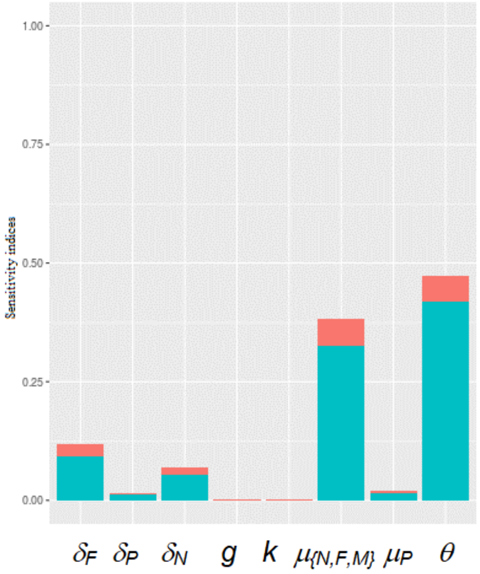

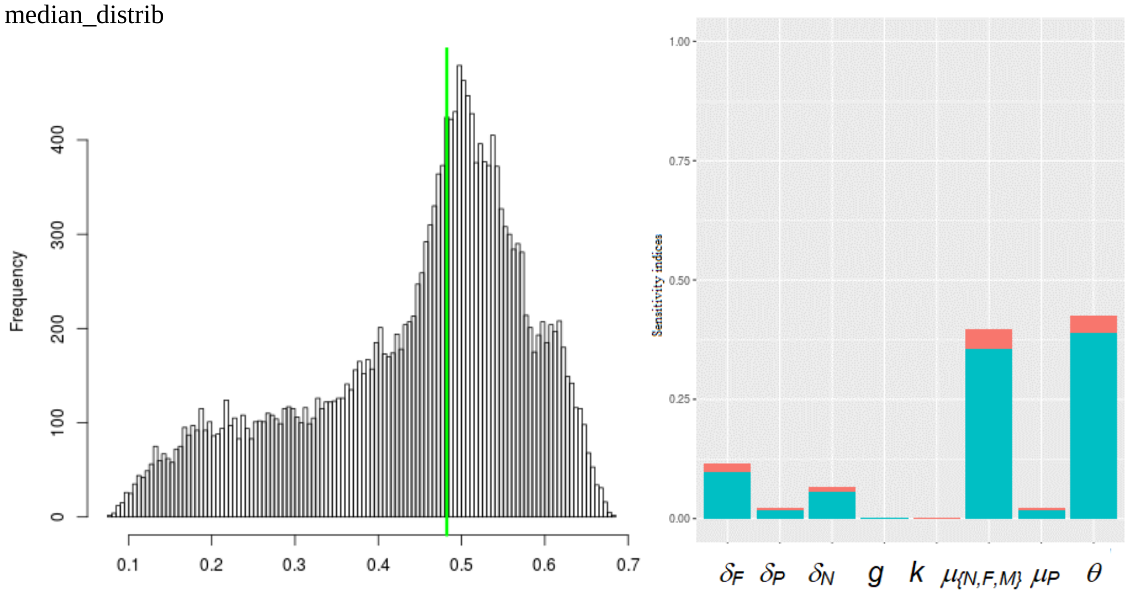

**Fig. S8.** Complementary results of the global sensitivity analysis. Left: output distributions. Right: principal effect (in blue) and interaction (in red) sensitivity indices for each output per varying parameter (refer to main text for parameter definition and to Table S3 for output definition). Results on popMean output are in the main text. Green line: no-control scenario.

#### 6. Additional information about control

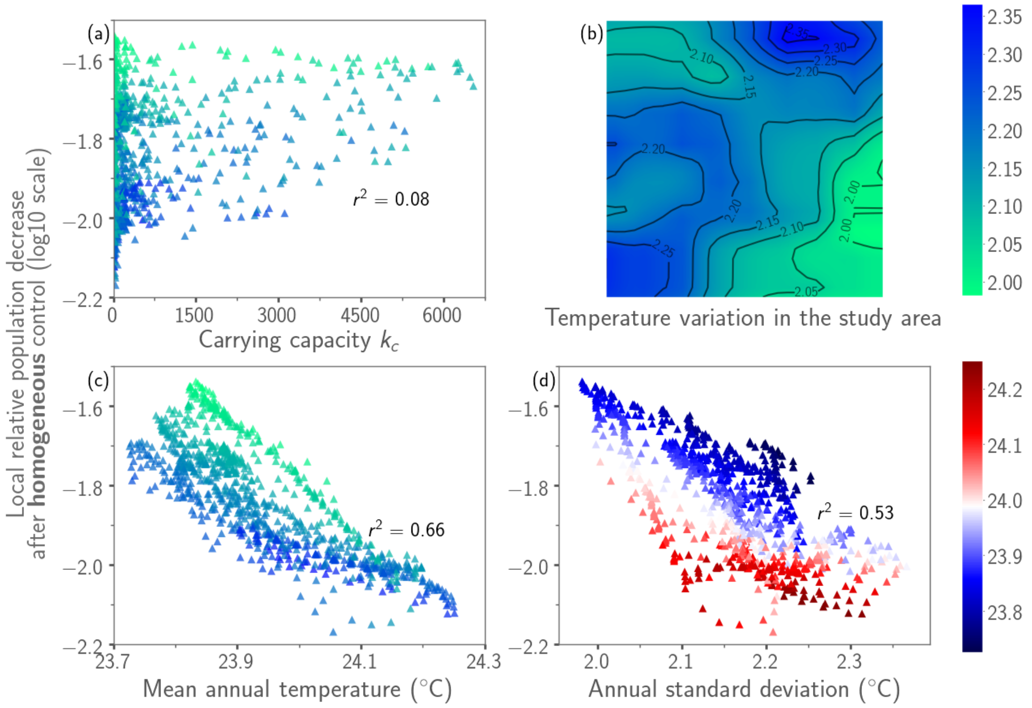

**Fig. S9.** Local efficacy of a homogeneous control applied in all pixels (female life expectancy of 35 days), with the lowest values of the relative population decrease (y-axis) representing the highest control efficacy. A: no correlation was observed with the local carrying capacity (A). B: spatial representation of the annual standard deviation of local temperatures (the colour bar was also used in A and C). C: the mean annual temperature was correlated with local efficacy. D: the annual standard deviation of temperature was correlated with local efficacy (points coloured by mean temperature to check the absence of correlation). Each point corresponds to a cell of the simulated grid.

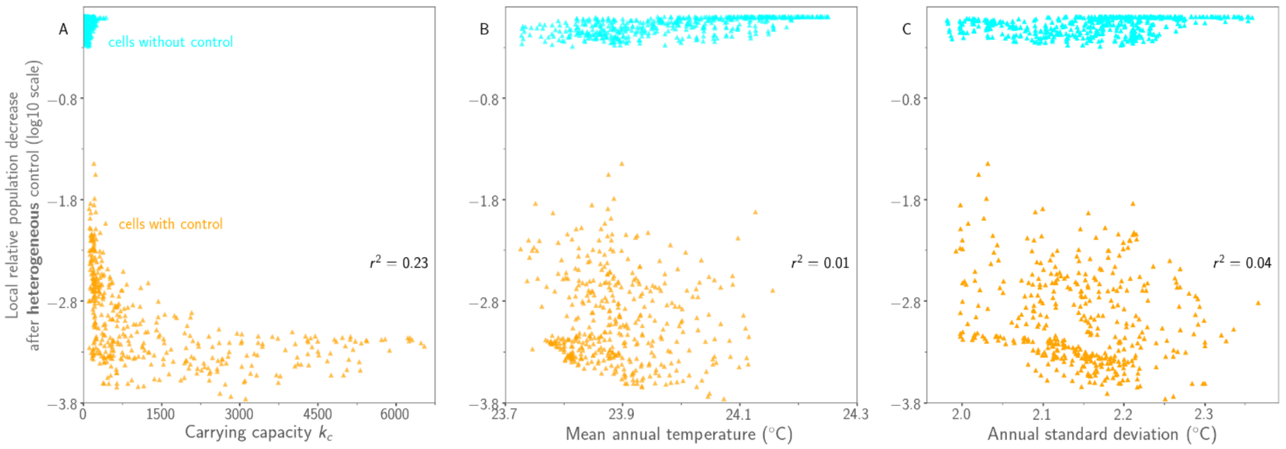

**Fig. S10.** Local efficacy of a heterogeneous control applied in 46% of the pixels (orange: treatment inducing a female life expectancy of 25 days; cyan: no treatment, female life expectancy of 60 days). No correlation was observed with local cell variables (A: carrying capacity; B: mean annual temperature; C: annual standard deviation of temperature).

#### 7. Code sources

To use the model code, go to: https://sourcesup.renater.fr/projects/spatial-tsetse/ or use directly the following command “git clone https://git.renater.fr/spatial-tsetse.git”
